# Spike mutation pipeline reveals the emergence of a more transmissible form of SARS-CoV-2

**DOI:** 10.1101/2020.04.29.069054

**Authors:** B Korber, WM Fischer, S Gnanakaran, H Yoon, J Theiler, W Abfalterer, B Foley, EE Giorgi, T Bhattacharya, MD Parker, DG Partridge, CM Evans, TM Freeman, TI de Silva, on behalf of the Sheffield COVID-19 Genomics Group, CC LaBranche, DC Montefiori

## Abstract

We have developed an analysis pipeline to facilitate real-time mutation tracking in SARS-CoV-2, focusing initially on the Spike (S) protein because it mediates infection of human cells and is the target of most vaccine strategies and antibody-based therapeutics. To date we have identified thirteen mutations in Spike that are accumulating. Mutations are considered in a broader phylogenetic context, geographically, and over time, to provide an early warning system to reveal mutations that may confer selective advantages in transmission or resistance to interventions. Each one is evaluated for evidence of positive selection, and the implications of the mutation are explored through structural modeling. The mutation Spike D614G is of urgent concern; it began spreading in Europe in early February, and when introduced to new regions it rapidly becomes the dominant form. Also, we present evidence of recombination between locally circulating strains, indicative of multiple strain infections. These finding have important implications for SARS-CoV-2 transmission, pathogenesis and immune interventions.

## Introduction

The past two decades have seen three major highly pathogenic zoonotic outbreaks of betacoronaviruses (Cui et al., 2019; de Wit et al., 2016; Fehr et al., 2017; Lu et al., 2020; Wu et al., 2020a). The first was Severe Acute Respiratory Syndrome Coronavirus (SARS-CoV) in 2002, which infected over 8,000 people and killed 800 (Graham and Baric, 2010). This was followed in 2012 by Middle East Respiratory Syndrome, MERS-CoV, a difficult to transmit but highly lethal virus, with 2,294 cases as of 2019, and 35% mortality (Cui et al., 2019; Graham and Baric, 2010). The third, SARS-CoV-2 is the cause of the severe respiratory disease COVID-19 (Gorbalenya et al., 2020). It was first reported in China in late December 2019 (Zhou et al., 2020), and triggered an epidemic (Wu et al., 2020b) that rapidly spread globally to become a pandemic of devastating impact, unparalleled in our lifetimes; today’s World Health Organization (WHO) situation report reads: over 2.5 million confirmed cases of COVID-19, and over 175,000 deaths (WHO Situation Report Number 94, April 23); tomorrow’s report will bring us new and markedly higher tallies of suffering, as the WHO continues to track the remarkable pace of expansion of this disease.

Three related factors combine to make this disease so dangerous: human beings have no direct immunological experience with this virus, leaving us vulnerable to infection and disease; it is highly transmissible; and it has a high mortality rate. Estimates for the basic reproductive ratio, R0, vary widely, but commonly range between 2.2-3.9 (Lv et al., 2020). Estimates of mortality, deaths per confirmed cases, also vary widely, and range between 0.5-15% (Zhou et al., 2020) (Mortality Analyses, John Hopkins University of Medicine). Differences in mortality estimates will reflect regional access to testing (a higher proportion of mild cases is detected when more testing is deployed), as well as regional differences in clinical care, and population differences in associated risk factors such as age. These basic numbers, R_0_ and mortality, are critical for public health response planning, but are difficult to resolve with confidence or to generalize across populations, given limited diagnostic testing and variations in the strategies of estimation.

Although the observed diversity among pandemic SARS-CoV-2 sequences is low, its rapid global spread provides the virus with ample opportunity for natural selection to act upon rare but favorable mutations. This is analogous to the case of influenza, where mutations slowly accumulate in the hemagglutinin protein during a flu season, and there is a complex interplay between mutations that can confer immune resistance to the virus, and the fitness landscape of the particular variant in which they arise (Wu et al., 2020c). Antigenic drift in influenza, the accumulation of mutations by the virus during an influenza season, provides the baseline variation needed to enable selection for antibody resistance across populations, and this drift is the primary reason we need to develop new influenza vaccines every few seasons. The seasonality of influenza is likely to be dictated in part by weather patterns (Chattopadhyay et al., 2018); longer seasonal epidemics allow selection pressure to continue over a more extended period, enhancing opportunities for the development of virus with novel antigenic surfaces that resist pre-existing immunity (Boni et al., 2006). SARS-CoV-2 is new to us; we do not yet know if it will wane seasonally as the weather warms and humidity increases, but our lack of pre-existing immunity and its high transmissibility relative to influenza are among the reasons it may not. If the pandemic fails to wane, this could exacerbate the potential for antigenic drift and the accumulation of immunologically relevant mutations in the population during the year or more it will take to deliver the first vaccine. Such a scenario is plausible, and by attending to this risk now, we may be able avert missing important evolutionary transitions in the virus that if ignored could ultimately limit the effectiveness of the first vaccines to clinical use.

There is clearly an urgent need to develop an effective vaccine against SARS-CoV-2, as well as antibody-based therapeutics (Kumar et al., 2020). Over 62 vaccine approaches are currently being explored, and a wide variety of candidate SARS-CoV-2 vaccines are in development (Landscape of COVID-19 Variants, WHO). Most of these vaccine approaches target the trimeric Spike protein (S) with the goal of eliciting protective neutralizing antibodies. Spike mediates binding and entry into host cells and is a major target of neutralizing antibodies (Chen et al., 2020; Yuan et al., 2020). Each Spike monomer consists of an N-terminal S1 domain and a membrane-proximal S2 domain, which mediate receptor binding and membrane fusion, respectively (Hoffmann et al., 2020; Walls et al., 2020; Wrapp et al., 2020). Notably, current immunogens and testing reagents are generally based on the Spike protein sequence from the index strain from Wuhan (Wang et al., 2020). SARS-CoV-2 is closely related to SARS-CoV; the two viruses share ~79% sequence identity (Lu et al., 2020) and both use angiotensin converting enzyme-2 (ACE2) as their cellular receptor (Hoffmann et al., 2020; Li et al., 2005; Wrapp et al., 2020), however the SARS-CoV-2 S-protein has a10-20-fold higher affinity for ACE2 than the corresponding S-protein of SARS-CoV (Wrapp et al., 2020). It remains to be seen to what extent lessons learned from SARS-CoV are helpful in formulating hypotheses about SARS-CoV-2, but SARS-CoV studies suggest that the nature of the antibody responses to the Spike protein are complex. In SARS-CoV infection, neutralizing Abs are generally thought to be protective; however, rapid and high neutralizing Ab responses that decline early are associated with greater disease severity and a higher risk of death (Ho et al., 2005; Liu et al., 2006; Temperton et al., 2005; Zhang et al., 2006). Furthermore, some antibodies against Spike mediate antibody dependent enhancement (ADE) of SARS-CoV (Jaume et al., 2011; Kam et al., 2007; Wan et al., 2020; Wang et al., 2014; Yilla et al., 2005; Yip et al., 2016; Yip et al., 2014). Because of the short duration of the outbreak, there were no efficacy trials of SARS-CoV vaccines and therefore we lack critical information that would help guide SARS-CoV-2 vaccine development.

Given Spike’s vital importance both in terms of viral infectivity and as an antibody target, we felt an urgent need for an “early warning” pipeline to evaluate Spike pandemic evolution. Our primary intent is to identify dynamically changing patterns of mutation indicative of positive selection for Spike variants. Also, because recombination is an important aspect of coronavirus evolution (Graham and Baric, 2010; Li et al., 2020; Rehman et al., 2020), we also set out to determine if whether recombination is playing a role in SARS-CoV-2 pandemic evolution. Here we describe a three-stage data pipeline (analysis of GISAID data, structural modeling of sites of interest, and experimental evaluation) and the identification of several sites of positive selection, including one (D614G) that may have originated either in China or Europe, but begin to spread rapidly first in Europe, and then in other parts of the world, and which is now the dominant pandemic form in many countries.

## Results

### Overview

Over the past two months, the HIV database team at Los Alamos National Laboratory has turned to developing an analysis pipeline to track in real time the evolution of the SARS-CoV-2 Spike (S) protein in the COVID-19 pandemic, using the Global Initiative for Sharing All Influenza Data GISAID SARS-CoV-2 sequence database as our baseline (Sup. Item 1 is the GISAID acknowledgments table, listing all the groups who contribute sequences to this global effort) (Elbe and Buckland-Merrett, 2017; Shu and McCauley, 2017). GISAID is the primary SARS-CoV-2 sequence database resource, and our intent is to complement what they provide with visualizations and summary data specifically intended to support the immunology and vaccine communities, and to alert the broader community to changes in frequency of mutations that might signal positive selection and a change in either viral phenotype or antigenicity. New global sequences arrive at GISAID at a furious pace; currently, hundreds of new SARS-CoV-2 sequences are added each day (Sup. Fig 1). By automating a series of key analysis steps with frequent inputs of GISAID data, critical analysis is provided in real time. The figures for this paper were based on an April 13, 2020 download of the GISAID data to enable the preparation of this manuscript. Our analysis pipeline (www.cov.lanl.gov) enables readers to reproduce the key figures (*e.g.* Figs. 1-3, and an updated table of sites of interest, Item S2) based on contemporary data downloaded from GIASID. The pipeline was developed in collaboration with the neutralizing antibody evaluation team at Duke University, who are in the process of creating a neutralizing antibody pseudovirus testing facility to experimentally resolve the virological and immunological implications of mutations of interest in Spike. While developing the computational tools to enable the analysis pipeline, the Los Alamos group has concurrently provided reports, at roughly two-week intervals, to colleagues at Duke as well as to others in the community who are working in the early design and testing phases of Spike-targeting vaccines and immunotherapeutics. This paper is essentially a formalized version of the fourth such report, based on the April 13^th^ GISAID data, and marks the transition of our analysis pipeline to a public resource and webpage (www.cov.lanl.gov).

**Fig. 1.**
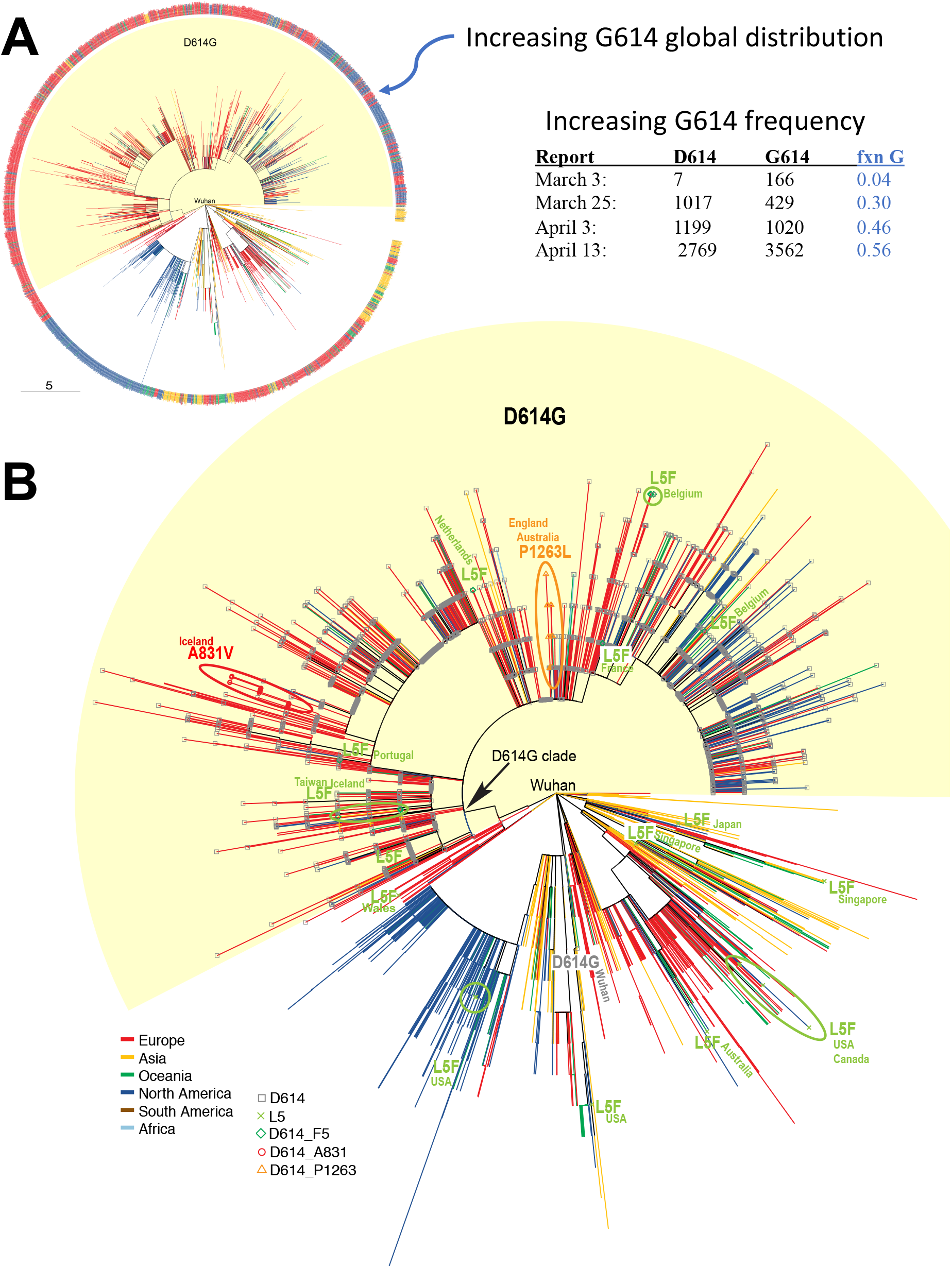
Phylogenetic Trees based on 4,535 trimmed full genome SARS C0V02 alignments from GISAID. **A)** A basic neighbor joining tree, centered on the Wuhan reference strain, with the GISAID G clade (named for the D614G mutation, though a total of 3 base changes define the clade) are highlighted in yellow. The regions of the world where sequences were sampled are indicated by colors. By early April, G614 was more common than the original D614 form isolated from Wuhan, and rather than being restricted to Europe (red) it had begun to spread globally. **B)** The same tree expanded to show interesting patterns of Spike mutations that we are tracking against the backdrop of the phylogenetic tree based on the full genome. Note two distinct patterns: mutations that predominantly appear to be part of a single lineage (P1263L, orange in the UK and Australia, and also A831V, red, in Iceland), versus a mutation that is found in very different regions both geographically and in the phylogeny, indicating the same mutation may be independently arising and sampled (L5F green, rare but found in scattered locations worldwide). A chart showing how GISAID sequence submissions increase daily is provided in Fig. S1. The tree shown here can be recreated with contemporary data downloaded from GISAID at www.cov.lanl.gov. The tree shown here was created using PAUP (Swofford, 2003); the trees generated for the website pipeline updates are based on parsimony (Goloboff, 2014).

Our analysis pipeline begins by downloading the GISAID data, then discarding partial and problematic sequences using stringent inclusion criteria, and then trimming jagged ends of the sequences back to the beginning of the first open reading frame (orf) and the end of the last orf. We use this to create two basic codon-aligned alignments (Kurtz et al., 2004): a more comprehensive Spike alignment (6,346 sequences as of April 13^th^), to monitor mutations that are beginning to accrue in Spike while maximizing the sample size; and a full genome alignment (4,535 sequences as of April 13^th^), to enable tracking the Spike mutations in a phylogenetic context informed by the evolutionary trajectories of the full genome (Fig. 1). Frequent public updates of analyses based on these alignments are provided through the pipeline.

Mutations among the pandemic SARS-CoV-2 sequences are sparse, limiting the applicability of traditional sequence-based methods for detecting positive selection. An alternative analysis framework for identifying positive selection can be taken, however, based on GISAID data, which provides a rich database of thousands of sequences linked to geographic information and sampling dates. This enables the tracking of sites for early indications of positive selection by identifying shifts in mutational frequencies over time. Early indicators include: (i) an increasing frequency of sequences that exhibit a particular mutation over time in a local region; (ii) frequent recurrent identification of a particular mutation in different geographic regions, and in different regions of the phylogenetic tree; (iii) the use of different codons to encode the same recurrent amino acid; and (iv) tight clusters of mutations in linear or structural sequence space. Because Spike mutations are rare, we set a low threshold for a site to be deemed “of interest” for further tracking. Thus, when a mutation is found in 0.3% of the sequences, we begin to track it by exploring its evolutionary trajectory and modeling its structural implications, *e.g*., its potential impact on antibody binding sites, trimer stability, and glycosylation patterns. Accumulation of mutations in known antibody epitopes are a special focus, including sites in or near the Spike Receptor Binding Domain (RBD), as well as sites that are near an immunodominant enhancing antibody epitope from the first SARS epidemic that centers on the mutation D614G (Wang et al., 2016); thus, a lower tracking threshold of 0.1% has been adopted for sites in Spike that are within 4 Å of these epitopes in the Spike 3D structure. If, given these criteria, a site merits further characterization, the experimental pipeline is triggered, and reagents are ordered and put into the queue for experimental evaluation of its impact on infectivity, antigenicity, neutralization sensitivity, and capacity to bind to the ACE2 receptor.

A summary of the sites of interest (as of April 13^th^) included 13 positions, as well as one local cluster of mutations (Table 1); a detailed report for each site is provided in supplemental data Item S2. Some of the sites are diminishing in frequency, and may have simply reflected a local sampling artifact from an earlier data set; they are still included in Item S2 because they reached the 0.3% threshold in a past sample of GISAID. Still others are increasing in frequency or persisting (Sup. Item 2), or have other interesting features suggesting they merit continued monitoring, such as recurrence in many regions of a phylogenetic tree (Fig. 1), or being located in a part of Spike that is of structural interest. We discuss a Spike mutation of particular interest, D614G, at some length, and then briefly summarize the other sites of interest. D614G is increasing in frequency at an alarming rate, indicating a fitness advantage relative to the original Wuhan strain that enables more rapid spread.

**Table 1.**
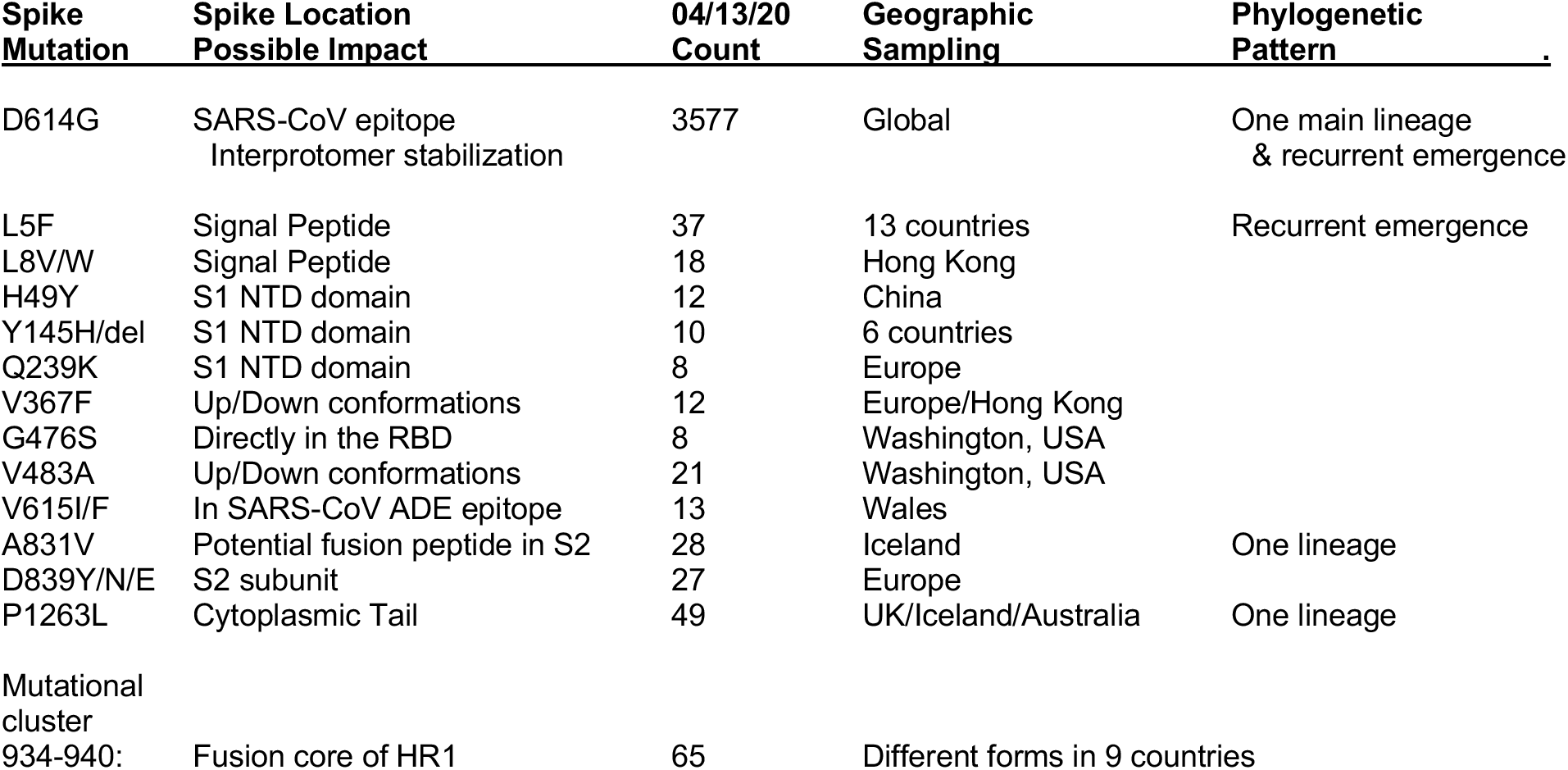
Table summarizing the mutations we are following in Spike. With the exception of D614G, all other mutations in Spike remain rare; we will nonetheless monitor them for potential immunological impact and/or for increased frequencies regionally or globally as the pandemic progresses. The NTD is the N-terminal domain, S2 is a membrane fusion subunit, and HR1 the first heptad repeat region (Lv et al., 2020). Up/Down conformations refer to a change in state in which the up conformation exposes the RBD (Kirchdoerfer et al., 2016; Kirchdoerfer et al., 2018). The SARS-CoV epitope was identified from the first SARS epidemic, and is the immunodominant linear antibody epitope observed in natural infection and in animal models. For the geographic sampling, we only list one country or region if it dominates a sample. For details see the complete listing of sequences with given mutation in Item S1.

### The D614G mutation

#### Increasing frequency and global distribution

The mutation D614G (a G-to-A base change at position 23,403 in the Wuhan reference strain) was the only site of interest identified in our first Spike mutation report in early March (Fig. 1A); it was found 7 times in 183 sequences that were available at the time. Four of these seven first D614G strains were sampled in Europe, and one each in Mexico, Brazil, and in Wuhan. In 5/7 cases, D614G was accompanied by 2 other mutations: a silent C-to-T mutation in the nsp3 gene at position 3,037, and a C-to-T mutation at position 14,409 which results in an RNA-dependent RNA polymerase (RdRp) amino acid change (RdRp P323L).The combination of these three mutations forms the basis for the clade that soon emerged in Europe (Fig. 1). By the time of our second report in mid-March, D614G was being tracked at GISAID due to its high frequency, and referred to as the “G” clade; it was present in 29% of the global samples, but was still found almost exclusively in Europe. It was recently reported by Pachetti *et al.* to be found in Europe and absent from regions globally, presumably because they capture GISAID data in roughly the time frame as our second report (Pachetti et al., 2020). The data available for study mid-March, given an approximate 2-week lag time between sampling and reporting, were consistent with the possibility of a founder effect in Europe resulting in spread across the continent, coupled with an increase in European sampling in the database. However, an early April sampling of the data from GISAID showed that G614’s frequency was increasing at an alarming pace throughout March, and it was clearly showing an ever-broadening geographic spread (Fig. 1A).

To differentiate between founder effects and a selective advantage driving the increasing frequency of the G clade in the GISAID data, we applied the suite of tools that we had been developing for the SARS-CoV-2 analysis pipeline (Fig. 2, Fig. 3, Fig. S2 and Fig. S3). A clear and consistent pattern was observed in almost every place where adequate sampling was available. In most countries and states where the COVID-19 epidemic was initiated and where sequences were sampled prior to March 1, the D614 form was the dominant local form early in the epidemic (orange in Figs. 2 and 3). Wherever G614 entered a population, a rapid rise in its frequency followed, and in many cases G614 became the dominant local form in a matter of only a few weeks (Fig. 3 and S3).

**Fig. 2.**
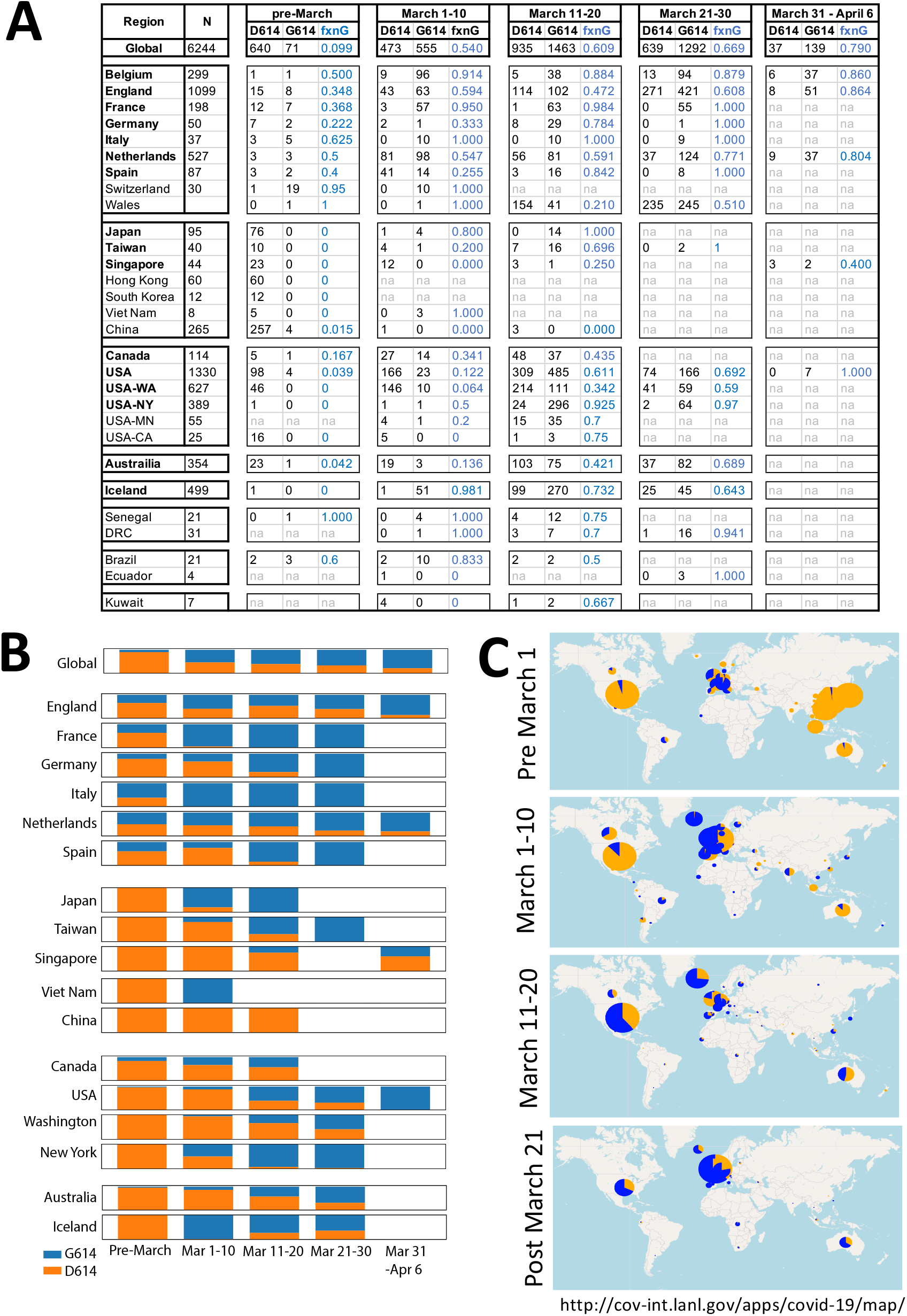
The proportion of sequences carrying the D614G mutation is increasing in every region that was well sampled in the GISAID database through the month of March. **A)** A table showing the tallies of each form, D614 and G614 in different countries and regions, starting with samples collected prior to March 1, then following in 10 day intervals. **B)** Bar charts illustrating the relative frequencies of the original Wuhan form (D614, orange), and the form that first emerged in Europe (G614, blue) based on the numbers in part (A). A variation of this figure showing actual tallies rather than frequencies, so the height of the bars represent the sample size, is provided as Fig. S2. **C)** A global mapping of the two forms illustrated by pie charts over the same periods. The size of the circle represents sampling. An interactive version of this map of the April 13^th^ data, allowing one to change scale and drill down in to specific regions of the world is available at https://cov.lanl.gov/apps/covid-19/map, and updates of this map based on contemporary data from GISAID are provided at www.cov.lanl.gov.

**Fig. 3.**
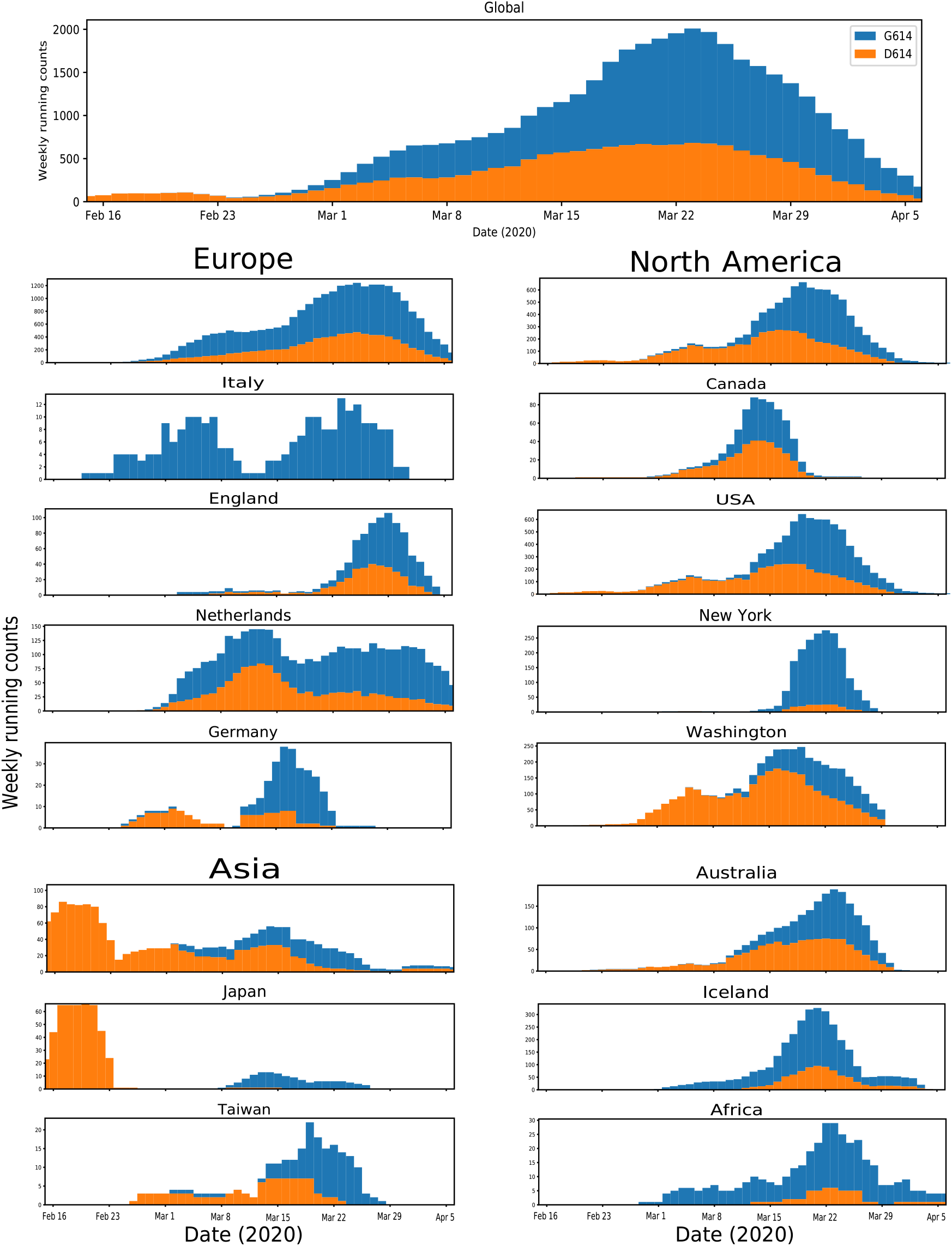
Running weekly average counts showing the relative amount of D614 (orange) and G614, (blue) in different regions of the world. In almost every case soon after G614 enters a region, it begins to dominate the sample. Fig. S3 shows the same data, illustrated as a daily cumulative plot. Plots were generated with Python Matplotlib (Hunter, 2007). The plots shown here and in Fig. S3 can be recreated with contemporary data from GISAID at www.cov.lanl.gov.

In Europe, where the G614 first began its expansion, the D614 and G614 forms were co-circulating early in the epidemic, with D614 more common in most sampled countries, the exceptions being Italy and Switzerland (Fig. 2A). Through March, G614 became increasingly common throughout Europe, and by April it dominated contemporary sampling (Fig. 2 and 3). In North America, infections were initiated and established across the continent by the original D614 form, but in early March, the G614 was introduced into both Canada and the USA, and by the end of March it had become the dominant form in both nations. Washington state, the state with the greatest number of available GISAID SARS-CoV-2 sequences from the USA, exemplifies this pattern (Figure 3 and S3), and a similar shift over time is evident in many other states with samples available throughout March (detailed data by state provided in Item S2). Sequences from New York were poorly sampled until some days into March (Fig. 3), and the G614 form was the predominant form; the G614 form was coming into prominence elsewhere in the USA by that time. Thus it is not clear whether the local pandemic was seeded by European contacts, as suggested in (Brufsky, 2020), if it was seeded by contacts from within the USA where it had already achieved high prevalence, or by a combination of both routes. Australia follows the same transition pattern, from D614 to G614 dominance, as the USA and Canada. Iceland is the single exception to the pattern; there sequencing was extensive and the epidemic seems to have started with a G614 form, but there was a transient increase in the D614 form, which then persisted at constant low level (Fig. 3 and S3). Asian samples were completely dominated by the original Wuhan D614 form through mid-March, but by mid-March in Asian countries outside of China, the G614 form was clearly established and expanding (Figs 2 and 3). The status of the D614G mutation in China remains unclear, as very few Chinese sequences in GISAID were sampled after March 1. South America and Africa remain sparsely sampled, but are shown for completeness in Figs. 2 and 3; for details see Item S2. Taken together, the data shows that G614 confers a selective advantage that is repeatedly reflected by dramatic shifts to G614 forms in regional epidemics over a period of several weeks.

The earliest D614G mutation in Europe was identified in Germany (EPI_ISL_406862, sampled 1/28/2020), and it was accompanied by the C-to-T mutation at 3,037, but not by the mutation at 14,409. Of potential interest for understanding the origin of the G614 clade, the G614 form of the virus was also found 4 times in China among early samples (Item S2). A Wuhan sequence (EPI_ISL_412982) sampled on 2/7/2020 had the D614G mutation, but it did not have either of the two accompanying mutations, and this single D614G mutation may have arisen independently (Fig. 1). The other three cases were potentially related to the German sequence. One sequence sampled on 1/24/2020 was from Zhejiang (EPI_ISL_422425), which borders Shanghai, and it had all three mutations associated with the G clade that expanded in Europe. The other two samples were both from Shanghai, and sampled on 1/28/2020 and 2/6/2020 (EPI_ISL_416327, EPI_ISL_416334); like the German sequence they lacked the mutation at 14,409. Given that these early Chinese sequences were also highly related to the German sequence throughout their genomes, it is possible that the G614 may have originated either in China or in Europe as it was present in both places in late January. There are no recent GISAID sequences from Shanghai or Zhejiang at this time, so we do not know if G614 was preferentially transmitted there, but the D614 form was prevalent in Shanghai in January and early February.

#### D614G and potential mechanisms for enhanced fitness

There are two distinct conceptual frameworks that may explain why the D614G mutation is associated with increased transmission. The first is based on structure. D614 is located on the surface of the spike protein protomer, where it can form contacts with the neighboring protomer. Examination of the cryo-EM structure (Wrapp et al., 2020) indicates that the sidechain of D614 potentially can form a hydrogen bond with T859 of the neighboring protomer as shown in Fig. 4C; the strength of Asp-Thr hydrogen bonding has been well documented (Kandori et al., 2001). This protomer-protomer hydrogen bonding may be of critical importance as it can bring together a residue from the S1 unit of one protomer to the S2 unit of the other protomer Fig. 4D. These two sites in the spike protein bracket both the dibasic furin- and S2-cleavage sites; thus, it is possible that the *D614G* mutation diminishes the interaction between the S1 and S2 units, facilitating the shedding of S1 from viral-membrane-bound S2. An alternative structure-based hypothesis is that this mutation may impact RBD-ACE2 binding (albeit indirectly since this site is not proximal to the binding interface). The RBD needs to be in the “up” position to engage with the ACE2 receptor, and it is possible that this site could allosterically alter the transitions of “up” and “down”. In the only known SARS-CoV-2 Spike structure with all RBDs in “down” position, the distances between *D614* and T859 remain the same between all protomers (Walls et al., 2020). However, in the structures with one RBD “up,” the distances between these residues are altered (Walls et al., 2020; Wrapp et al., 2020). The distance between the “up” protomer and the “down” neighboring protomer (clockwise) is slightly longer than the rest of the protomer-protomer distances (Fig. S4D). These slight changes are, however, most likely within the conformational fluctuations of a dynamic spike trimer. More detailed experimental and modeling studies are needed to elucidate the effect of this mutation on RBD transitions.

**Fig. 4.**
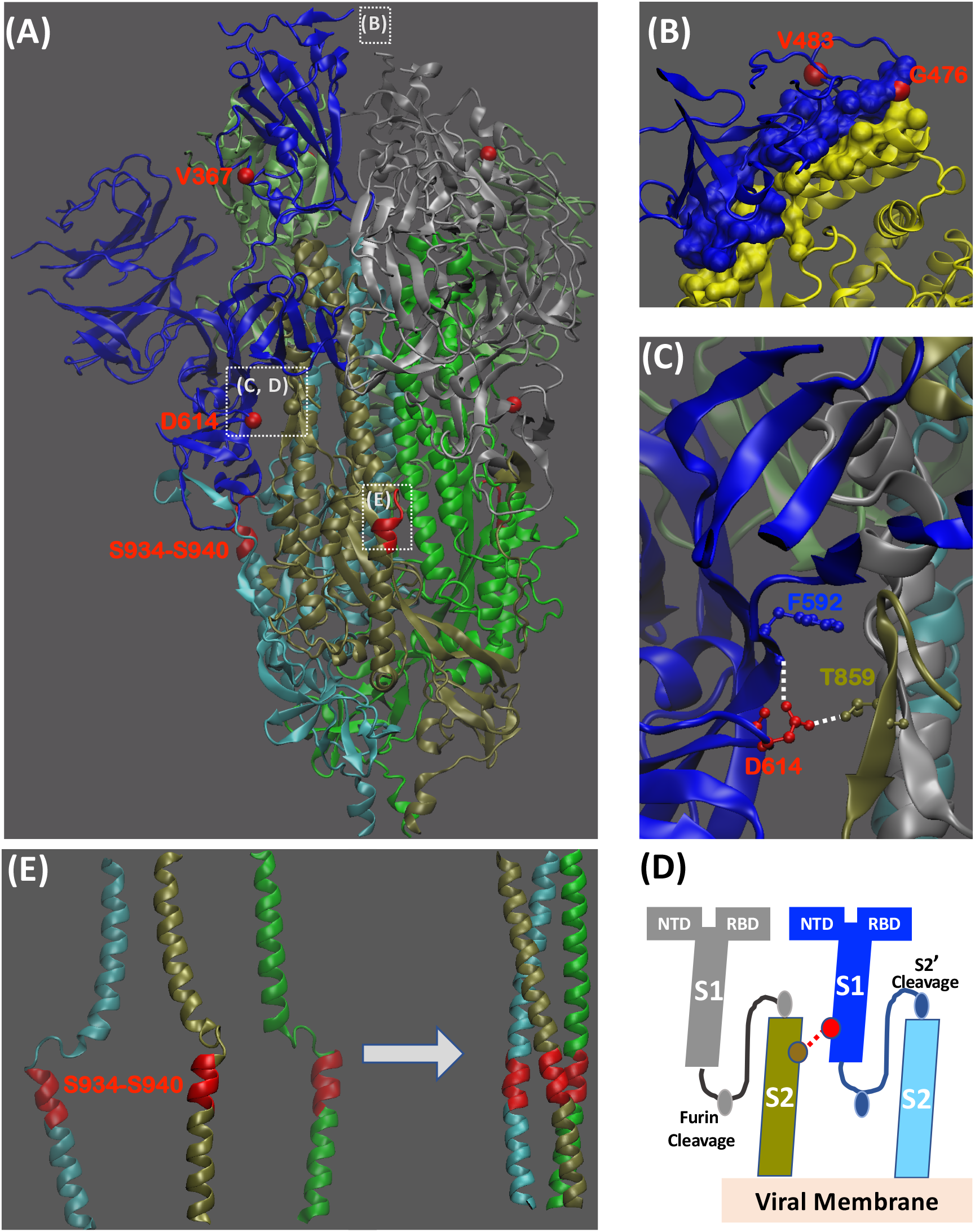
Structural mapping key mutational sites in the Spike protein. **A)** Mutational sites span S1 and S2 structural units of the Spike protein (PDB:6VSB). Different colors are used to distinguish the protomers and sub-units. S1 and S2 sub-units are defined based on the furin cleavage site (protomer #1: S1-blue, S2-cyan, protomer #2: S1-grey, S2-tan, protomer #3: S1-light green, S2-dark green). The RBD of protomer #1 is in “UP” position for engagement with ACE2 receptor. Red color is used to indicate individual mutational sites (ball). The dashed squares with labels indicate forthcoming detailed investigations in subsequent images. **B)** Mutational sites near the RBD (blue)-ACE2 (yellow) binding interface. The interfacial region is shown as a surface (PDB: 6M17). **C)** The proximity of D614 to T859 from the neighboring protomer. The white dashed lines indicate the possibility for forming hydrogen bonds. **D)** A cartoon is used to capture how the potential protomer-protomer interactions shown in (C) brings together D614 from S1 unit of one protomer to the T859 from S2 unit of the neighboring protomer. **E)** Cluster of mutations, *S934-S940*, in the HR1 region of the Spike protein. These residues occur in a region that undergoes conformational transition during fusion. The left and right images show the pre-fusion (PDB:6VSB) and post-fusion (PDB: 6LXT) conformations of this HR1 region. Structural implications of different sites are noted in Fig. S4. Structural evaluations and rendering of three dimensional images were carried out using Visual Molecular Dynamics (VMD) (Humphrey et al., 1996).

The second way the D614G mutation might impact transmission is immunological. D614 is embedded in an immunodominant linear epitope in the original SARS-CoV Spike, S597–603. This peptide had a very high level of serological reactivity (64%), and induced long term B-cell memory responses in convalescent-phase sera from individuals infected during the original SARS-CoV 2002 epidemic (Wang et al., 2016). Antibodies against this peptide mediate antibody-dependent enhancement (ADE) of SARS-CoV infection by an epitope-sequence-dependent mechanism, both *in vitro*, and *in vivo* in rhesus macaques (Wang et al., 2016). The minimal linear core epitope for the ADE mediating antibodies in SARS-CoV was *LYQDVNC* (SARS-CoV S_597-603_), and this peptide is immediately proximal to a peptide that is targeted by potentially beneficial neutralizing antibodies (SARS-CoV S_604-625_). The ADE target peptide spans the SARS-CoV-2 D614 site, and is identical to the equivalent region in SARS-CoV, S_611-617_. Wang et al. noted proximity of this epitope to the RBD, and speculated that antibody binding may mediate a conformational change in Spike that increases RBD-ACE2 interaction resulting in the enhancing effect. This mechanism is notably different from the more common Fc receptor-mediated mechanism of SARS-CoV ADE reported by others, and which occurs in both the presence and absence of ACE2 (Jaume et al., 2011; Kam et al., 2007; Wan et al., 2020; Wang et al., 2014; Yilla et al., 2005; Yip et al., 2016; Yip et al., 2014). Thus, based on currently available information, there are several ways the D614G mutation may impact Spike’s infectivity: it may improve receptor binding, fusion activation, or ADE antibody elicitation. Another mechanism for the shift to the G614 form at later times points might simply be through antigenic drift mediating antibody escape. If the D614G mutation in SARS-CoV-2 was impacting neutralizing antibody sensitivity as well as, or instead of, the ADE activity observed in the SARS-CoV study, D614G could also be mediating escape that makes individuals susceptible to a second infection.

#### D614G and clinical outcome

We were concerned that if the D614G mutation can increase transmissibility, it might also impact severity of disease. Because clinical outcome data are not available in GISAID, we focused on a single geographic region, Sheffield, England, where a large data set existed and was made available for an initial exploration of this question. SARS-CoV-2 sequences were generated from 453 individuals presenting with COVID-19 disease at the Sheffield Teaching Hospitals NHS Foundation Trust. Sheffield followed the pattern observed through much of world, starting out with D614, and shifting to predominantly G614 by the end of March. The Sheffield data included age, gender, date of sampling, cycle threshold (CT) for positive signal in E gene-based RT-PCR (used here as a surrogate for relative levels of viremia) (Corman et al., 2020), and clinical status: outpatient (OP), inpatient (IP, requiring hospitalization), or admittance into the intensive care unit (ICU). Because the numbers admitted into in the ICU were small, and because that information was not readily available for all subjects, we grouped the IP and ICU together for most of the statistical analysis. As anticipated, there was a significant relationship between hospitalization and age (Fig. S5) (Wilcoxon p < 2.2e-16, median age 74, interquartile range (IR) 58-83 for the hospitalized patients, versus a median of age 43 (with IR 31-53) for the outpatients (Dowd et al., 2020; Promislow, 2020). Also, males were hospitalized more often than females (Fisher’s p = 1.95e-13) (Conti and Younes, 2020; Promislow, 2020). Furthermore, fewer PCR cycles were required for detection of virus among individuals who admitted to the hospital, indicating higher viral loads (Fig. S5) (Wilcoxon p = 4.8e-06, median 25, IR 21-28, versus median 19, IR 21-25) (Fig. S5) (Liu et al., 2020). *There was, however, no significant correlation found between D614G status and hospitalization status*; although the G614 mutation was slightly enriched among the ICU subjects, this was not statistically significant (Fig. 5C).

**Fig. 5.**
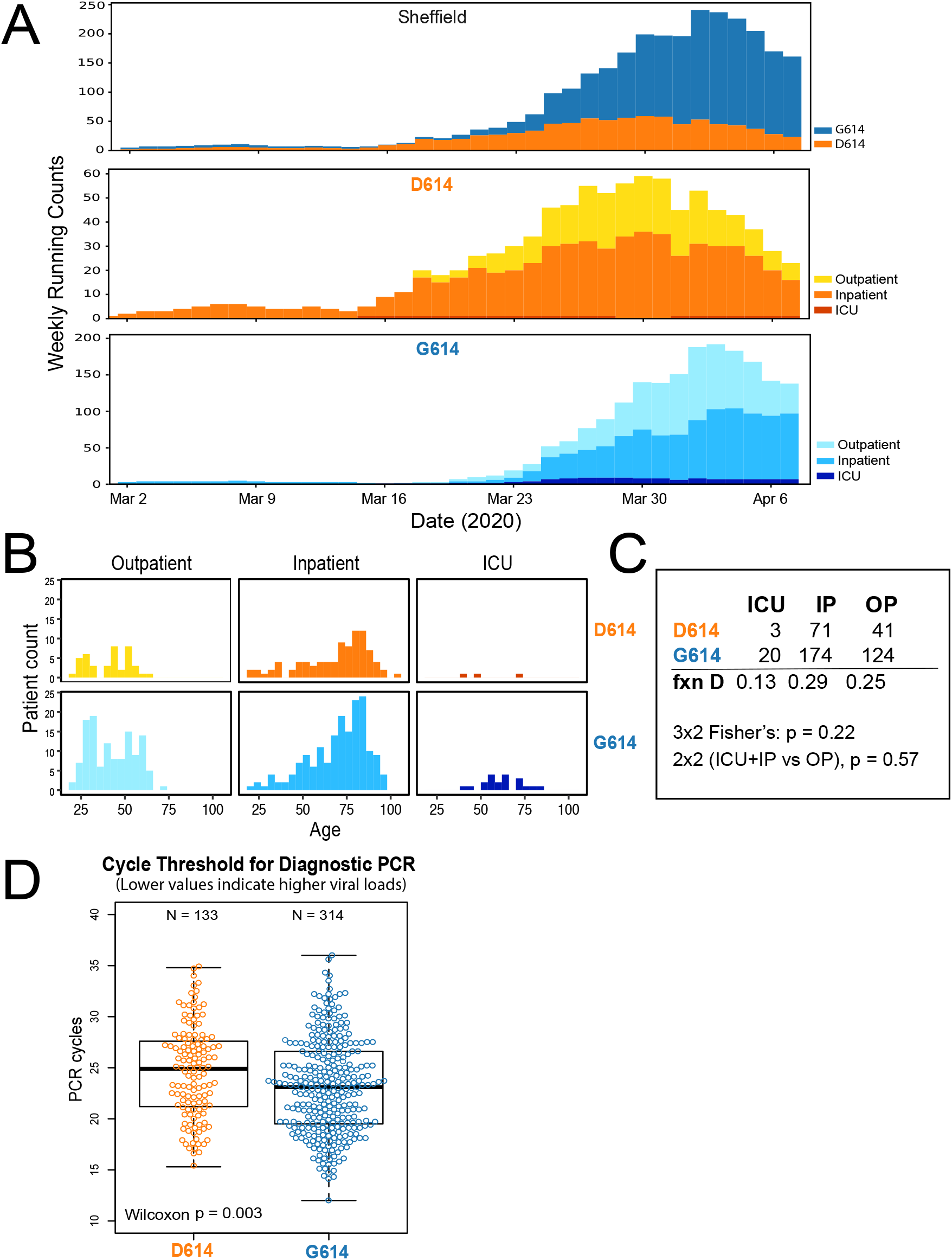
A comparison of the two forms of S D614G to clinical outcomes in 447 subjects with COVID-19, over the time span of the epidemic in Sheffield, England. **A)** Tracking the epidemic forms in Sheffield. The top panel shows that the course of the epidemic in Sheffield is the same as the course we documented throughout the globe (see Fig. 3), with G614 overtaking D614 as the dominant form of SARS-CoV-2. We were concerned that the rate of hospitalization might have varied over this time period, which could have biased the sample as G614 tended to be sampled later. The D614 and G614 panels show that the rate of hospitalized individuals from whom sequences were obtained, averaged per week, remained relatively constant across this time period for both groups. **B)** Age distribution between clinical status groups. We were also concerned that the age distribution of people visiting the hospital might have differed between the groups, as age is highly associated with high risk, but part **(B)** shows that this distribution is very similar, and there was no statistical difference between the groups overall (Wilcoxon p = 0.77, D614 had a median age of 60 (IR 44-80), while G614 had a median age of 59 (IR 43-77). **C)** D614G status was not statistically associated with hospitalization status. **D)** G614 was associated with fewer rounds of PCR required for detection, suggesting that people with a G614 virus had higher viral loads. Other associations with hospital status are shown in Fig. S5.

We were concerned sampling issues might have introduced a bias, particularly as the D614 viruses were more heavily sampled early, and G614 at later times, and clinical practice might have changed over the course of the time period. Age, however, was evenly distributed between the G614 and D614 hospitalized groups (Fig 5B), and the relative number of hospitalizations stayed constant throughout the study period in both the D614 and G614 groups (Fig 5A). We also performed a multivariate Generalized Linear Model (GLM) analysis, (Bates et al., 2015) with outpatient vs. hospitalized status as the outcome, and age, gender, D614, and PCR CT as potential predictors. PCR CT was a significant predictor of disease severity, after adjusting for age and gender, but D614G status was not. Again, age was the most significant predictor (p<2×10^−16^), followed by gender (p=2 ×10^−7^) and then PCR CT (p=1.9×10^−6^). D614G status did not significantly contribute to modeling hospitalization as an outcome, but there was a marginally significant interaction with PCR CT (p=0.04).

While D614G did not predict hospitalization, there was a significant shift in cycle threshold to fewer PCR cycles being required for detection among the group that carried G614 relative to D614 (Fig. 5D). This indicated that patients carrying the G614 mutation had higher viral loads (Wilcoxon p = 0.003, median 23.1, IR 19.5-26.6, versus median 24.9, IR 21.2-27.6). This comparison is limited by uncertainty regarding the time from infection at which sample was taken, and by the fact that PCR is an indirect measure of viral load; still it is notable that despite these limitations, a significant difference was observed.

### Recombination among pandemic sequences

Knowing that recombination plays an important role in coronavirus evolution generally (Graham and Baric, 2010; Li et al., 2020; Rehman et al., 2020), the possibility that recombination might be also be contributing significantly to evolution in the current pandemic seemed plausible, but difficult to detect using standard methods given how little variation there is among pandemic strains. Recombination requires simultaneous infection of the same host with different viruses, and the two parental strains have to be distinctive enough to manifest in a detectable way in the recombined sequence.

To determine if potential recombination events could be identified in geographically regional data sets, we applied a computational method called RAPR (Song et al., 2018) that we had originally developed to explore the evolutionary role of within-patient HIV recombination in acute HIV infection—another situation of low viral diversity. RAPR enables the comparison of all triplets (sets of three sequences) in an alignment, and applies a run-length statistic to evaluate the possibility of recombination. We began this analysis with the set of SARS-CoV-2 sequences from Washington State, as it was particularly well sampled set and from a geographically local population where co-infection might occur. Using RAPR, we found several recombination candidates, and the two most significant examples of these are shown in (Fig. 6). To identify these cases, we did all possible comparisons of three sequences in the sample set, and we show raw p-values given for the run-length statistic; these p-values did not withstand a multiple correction for the run length statistic. The statistic, however, only considered whether the similarities to the parents are clustered as expected in a recombination event, without regard to the plausibility of alternate hypotheses like recurring mutations. Given the very low rate of mutation among the pandemic sequences, however, the patterns of shared sites shown in Fig. 6 seem highly unlikely to have arisen by serial spontaneous mutation, thus either recombination *in vivo*, or possibly recombination *in vitro* as a result of contamination during PCR, are a better explanations of the observed data. Furthermore, RAPR analysis indicated that one of the recombinant forms in Washington founded a lineage that continued to spread. To identify other geographic regions where recombination might be occurring among the sequences in the global data set, we next focused on sequences sampled from areas with co-circulation of both haplotypes of the D614G mutation, as defined by 3 mutational events (Fig. 1): the a C-to-T mutation at position 3,037, a C-to-T mutation at position 14,408, and the G-to-A base change at position 23,403 that gives rise to the D614G amino acid change in spike. The original Wuhan D614 form carried the bases ‘C-C-G’ in these positions, and the G614 form carries the bases ‘T-T-A’. Given that these positions were well spaced in the genome, and the two forms were co-circulating in many communities over the month of March, we reasoned that deviations from either the ‘C-C-G’ or the ‘T-T-A’ patterns would be indicative of recombination. Such deviations were rare, found only in 0.8% of our full genome alignment of 4,535 sequences, but with examples found in Belgium, Netherlands, Minnesota, Spain, Iceland, Latvia, NanChang, and Australia. Using RAPR to identify likely recombination among these sets we were indeed able to identify multiple recombination candidates; some of the most significant examples from Iceland and the Netherlands are also shown in Fig. 6.

**Fig. 6.**
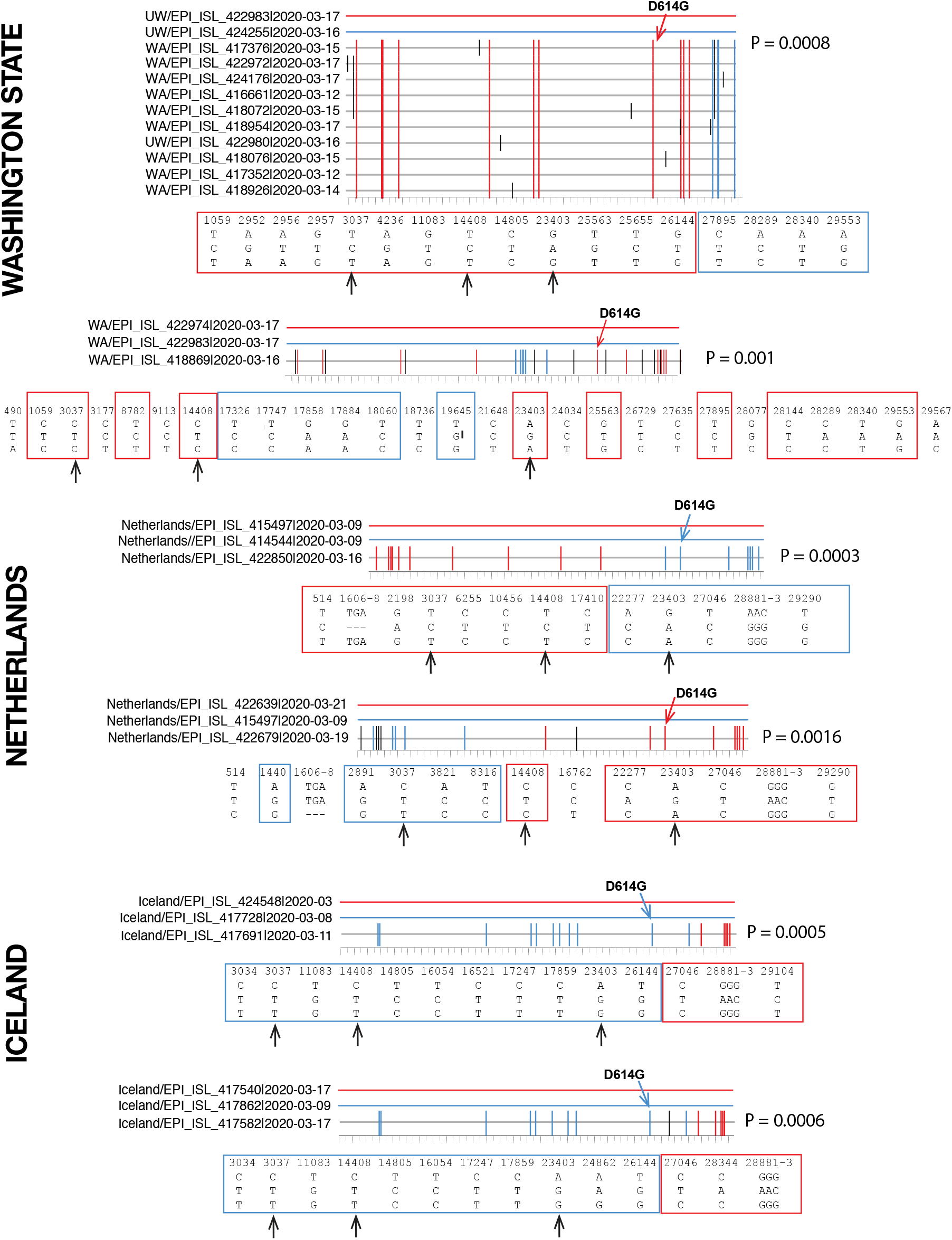
Representative examples of recombination detected in three different regions. Each graph illustrates a recombination event detected by the software RAPR. P-values are from the Wald–Wolfowitz runs test. While we are unlikely to have sampled actual recombination parents and child (the unique bases marked with black tics emphasize this), we have sampled representatives from each of the lineages that were likely to be involved in a recombination event. Putative parental strains are shown in red (top) and light blue (bottom), and the solid colored line represents the nearly 30,000 bases of their full length SAR-CoV-2 genomes. The recombinant child is shown below with color-coded tick marks representing mutations matching either parental strain, when the parents differ in the base at a given position. Nucleotides in black are distinct in the child and do not match either parent. Boxes below show nucleotides at each position of diversity across the triplet, with red or light blue boxes highlighting the parental strain they match. Black arrows show the D614G trio of mutations at positions 3037, 14408 and 23403 (D614G in the spike gene). The p-values are based on a run-length statistic, and are not corrected for multiple tests. **Top panel**: A recombination event in which the putative recombinant “child” gave rise to a cluster of 10 recombinants in the WA state sequence set. Below, a second example from the same sequence set shows a more complex recombination event with two distinct breakpoints. **Middle panel**: Two examples of recombination events detected in the sequence set from the Netherlands. **Bottom panel**: Two examples of recombination events detected in the sequence set from Iceland. In both of these cases the red and blue parents were similar, and breakpoints were similar, but the red and blue parents were distinct in a few positions. In cases from the Netherlands and Iceland, the haplotypes representing the GISAID G clade that carries the D614G mutation and the other two mutations were mixed, their positions are indicated with arrows.

### Additional sites of interest in Spike with accruing but rare mutations

#### Sites L5F and L8V

These are both signal peptide mutations, and it is difficult anticipate how they might impact the virus. Variation in the signal peptide of other viruses, for example HIV Env, can impact posttranslational modifications in the endoplasmic reticulum, including folding, expression levels, and glycosylation (Asmal et al., 2011; Upadhyay et al., 2018). The L5F mutation is intriguing because of its recurrence in many lineages throughout the SARS-CoV-2 phylogenetic tree, and in many different countries throughout the world (Fig 1). Once established, it is often regionally transmitted and contributes to multiple small local clusters (Fig. 1, Item S2) – *e.g.* a cluster of 5 infections in Iceland with identical in sequences all carrying 5F, and several comparable clusters in different states in the USA). This combination suggests that it may be a favorable mutation that tends to persist when it arises. Despite its recurrences, it has maintained but not increased in frequency, and continues to be found at roughly 0.6% of the global sampling through April.

L8V is potentially interesting because of a very different evolutionary trajectory. It is mostly found in a single lineage in Hong Kong, with one apparent recurrence in Canada. Because Hong Kong has not been recently sampled in GISAID, L8V’s overall global frequency appears to decline in April; however, L8V may in fact be increasing in frequency in Hong Kong over time and thus merits continuing scrutiny (Fig. S6)

#### Sites V367F, G476S, and V483A

There are three mutational sites, V367F, G476S, and V483A, that are found within the RBD domain (Fig. 4 A,B). Of these three, only G476S occurs directly at the binding interface of RBD and peptidase domain of ACE2. Fig. 4B shows that G476 is at the end of the binding interface that is predominantly driven by polar interactions (Yan et al., 2020). The closest ACE2 residue to G476 is Q24, which is near the end of α-1, the helix of ACE2 most engaged in RBD recognition. The ACE2 residue Q24 makes distinct electrostatic interactions with the backbone of A475, the neighboring residue to G476 (Fig. S4 B). The G476S mutation may contribute to the rigidification of the local loop region in RBD near the binding interface, or bring this flexible loop even closer to the interface. The mutation at site V483 does not directly contact ACE2 although it is on the same face of the RBD that forms the binding interface with the ACE2. Site V367 is at the opposite end from where RBD binds to ACE2 receptor; it is on the same face as the epitope of CR3022, a neutralizing antibody that was isolated from a convalescent SARS-CoV patient when at least two of the RBD regions of the Spike trimer are “up”, (Fig. S4 C) though no direct contacts between V367 and CR3022 are observed (Yuan et al., 2020).

Sites V367F and G476S were identified as mutations of interest in early smaller datasets, and appear to be diminishing in overall global frequency in later samples, but V367F merits continued scrutiny due to its potential for interactions with ACE2. The mutation V483A has predominately appeared in Washington State (Item S2). It is not increasing over time among samples from Washington, but it maintains a steady, albeit very low, presence (Fig. S6).

#### Sites H49Y, Y145H/del, Q239K

Each of these mutations are located in S1 N-terminal domain (NTD), a domain not well characterized functionally. The sites were identified as sites of interest in early smaller datasets, but appear to be diminishing in overall frequency in later samples. They recur in different countries, although K239 is predominantly found in the Netherlands.

#### Sites A831V and D839Y/N/E

A831V and D839 are both being maintained at ~0.4% in the global sample through April. A831V is found only in Iceland and is in a single lineage that is stable in frequency over time (Fig. S6), whereas D839 has been sampled in many countries in Europe. Both mutations occur in the region of fusion peptide, although neither position is structurally resolved in the cryo-EM reconstruction; site A831 is part of the canonical fusion peptide. We have also identified a small local cluster of mutations in S934-S940 (Fig. S7), focused in the fusion core of the HR1 (Xia et al., 2020), next to the region where the helix is broken in the trimeric pre-fusion spike (Table S2, **Fig. 4E**). This cluster is rich in Serine residues that have a high propensity to form hydrogen bonds. Upon spatially localizing in a helix with a motif SXSS (937-940) as seen here (937-943: SXSSTXS), it has the potential to enhance the association of helices. Previously, SXXSSXXT-like motifs have been shown to drive the association of trans-membrane helices (Dawson et al., 2002) and could be even more relevant given the association with amphipathic helices. Given that, this cluster in the HR1 region of S2 unit could impact conformational rearrangements as the S2 unit transitions from pre-fusion to post-fusion by enhancing the association of the HR1 trimer and maintaining amphipathicity when the a single HR1 helix is extended (Fig. S4). Even though such a motif is not seen in HIV gp41, it is observed near the MPER region in γ-retroviral glycoproteins and has been suggested as a possible conserved mechanism to drive oligomerization (Salamango and Johnson, 2015). (Of note, we had originally identified the **S943P** as a mutation of interest because it had met our threshold critia, and it was also located in the fusion core, but a closer examination of this mutation revealed that it was the result of a sequencing processing error (Freeman et al., 2020) (see Fig. S8, and methods for details).)

#### Site P1263L

This mutation is not included in the SARS-CoV-2 structure, but is near the end of the cytoplasmic tail of the Spike protein. The mutation is found mostly in the UK, in both England and Wales, but also in Australia, and is emerging as a single related lineage (Fig. 1). It is maintaining its frequency both globally as well as locally in the UK.

## Discussion

When we embarked on our SARS-CoV-2 analysis pipeline, our motivation was to identify mutations that might be of potential concern in the SARS-CoV-2 Spike protein as an early warning system for consideration as vaccine studies progress; we did not anticipate such dramatic results so early in the pandemic. In a setting of very low genetic diversity, traditional means of identification of positive selection have limited statistical power, but the incredibly rich GISAID data set provides an opportunity to look more deeply into the evolutionary relationships among the SARS-CoV-2 sequences in the context of time and geography. This approach revealed that viruses bearing the mutation Spike D614G are replacing the original Wuhan form of the virus rapidly and repeatedly across the globe (Fig. 2-3). We do not know what is driving this selective sweep, nor for that matter if it is indeed due the modified Spike and not one of the other two accompanying mutations that share the GISAID “G-clade” haplotype. The Spike D614G change, however, is consistent with several hypotheses regarding a fitness advantage that can be explored experimentally. D614 is embedded in an immunodominant antibody epitope, recognized by antibodies isolated from recovered individuals who were infected with the original SARS-CoV; this epitope is also targeted by vaccination in primate models (Wang et al., 2016). Thus, this mutation might be conferring resistance to *protective* D614-directed antibody responses in infected people, making them more susceptible to reinfection with the newer G614 form of the virus. Alternatively, the advantage might be related to the fact that D614 is embedded in an immunodominant ADE epitope of SARS-CoV (Wang et al., 2016), and perhaps the G614 form can facilitate ADE. Finally, the D614G mutation is predicted to destabilize inter-protomer S1-S2 subunit interactions in the trimer, and this may have direct consequences for the infectivity of the virus (Fig. 4). Increased infectivity would be consistent with rapid spread, and also the association of higher viral load with G614 that we observed in the clinical data from Sheffield, England (Fig. 5).

Many of the ways we anticipated we might find evidence of positive selection in Spike are being manifested among subset of sites with accruing mutations. While the D616G mutation is the only one that is dramatically increasing in frequency globally (Fig. 2-3), the L8V mutation may be on the rise in the local epidemic in Hong Kong (Fig. S7). To date, mutations are extremely rare in the Spike RBD, but the mutation G476S is directly in an ACE2 contact residue. The mutation L5F occurs in many geographic regions in many distinct clades, suggesting it repeatedly arose independently, and was selected to the extent that was frequent enough to be resampled, or is possibly a recurrent sequencing artifact. Finally, we have found evidence of recombination among regional sample sets (Fig. 6). Recombination among pandemic SARS-CoV-2 strains would not be not surprising, given that it is also found among more distant coronaviruses with higher diversity levels (Graham and Baric, 2010; Li et al., 2020; Rehman et al., 2020). Still, it has important implications. First, natural recombination cannot be detected without simultaneous coinfection of distinct viruses in one host. If the recombination events that are illustrated in Fig. 6 are indeed happening *in vivo*, co-infections that enable them might be happening prior to the adaptive immune response, or in series with reinfection occurring after the initial infection stimulated a response. Recombination may be more common in communities with less rigorous shelter-in-place and social distancing practices, in hospital wards with less stringent patient isolation because all patients are assumed to already be infected, or in geographic regions where antigenic drift has already begun to enable serial infection with more resistant forms of the viruses. Also, recombination provides an opportunity for the virus to bring together, into a single recombinant virus, multiple mutations that independently confer distinct fitness advantages but that were carried separately in the two parental strains.

Tracking mutations in Spike has been our primary focus to date because of the urgency with which vaccine and antibody therapy strategies are being developed; the interventions under development now cannot afford to miss their contemporary targets when they are eventually deployed. To this end, we built a data-analysis pipeline to explore the potential impact of mutations on SARS-CoV-2 sequences. The analysis is performed anew as the data becomes available through GISAID. Experimentalists can make use of the most current data available to best inform vaccine constructs, reagent tests, and experimental design. While the GISAID data used for the figures in this paper was frozen at April 13, 2020, many of the key figures included here are rebuilt on a frequent basis based on the newly available GISAID data. While our initial focus is on Spike, the tools we have developed can be extended to other proteins and mutations in subsequent versions of the pipeline. Meanwhile understanding both how the D614G mutation is overtaking the pandemic and how recombination is impacting the evolution of the virus will be important for informing choices about how best to respond in order to control epidemic spread and resurgence.

## Supporting information

Item S1: GISAID acknowledgement file April

## ACKNOWLEDGEMENTS

We thank Sir Andrew McMichael, Professor Sarah Rowland-Jones, and Dr. Xiao-Ning Xu for their invaluable help with bringing together the clinical and theoretical biology teams necessary for this study. Sequencing of SARS-CoV-2 samples was undertaken by the Sheffield COVID-19 Genomics Group as part of the COG-UK CONSORTIUM. COG-UK and supported by funding from the Medical Research Council (MRC) part of UK Research & Innovation (UKRI), the National Institute of Health Research (NIHR) and Genome Research Limited, operating as the Wellcome Sanger Institute. TIdS is supported by a Wellcome Trust Intermediate Clinical Fellowship (110058/Z/15/Z). Analyses strategies presented in this article were developed with the support of the Laboratory Directed Research and Development program of Los Alamos National Laboratory. Recombination analysis was conducted under project number 20200554ECR. The sequence data pipeline design and analysis of the structural immunological implications of Spike mutations was conducted under project number (20200706ER). The sequence data pipeline implementation was funded through the National Institute of Allergy and Infectious Diseases, National Institutes of Health, Department of Health and Human Services, under Interagency Agreement No. AAI12007-001-00000. We gratefully acknowledge the team at GISAID for creating the remarkable COVID-19 outbreak global database and resources, and the many authors from the oringiating and submitting laboratories of sequence data on which this analysis is based; see Item S1 is the acknowledgment table from GISAID at the time of our April 13^th^ data download, listing the many people responsible for generating the sequence data. Bless them.

## STAR METHODS

### RESOURCE AVAILABILITY

#### Lead Contact

Further information and requests for resources should be directed to and will be fulfilled by the Lead Contact, Bette Korber.

#### Materials Availability

This study did not generate new unique reagents.

#### Data and Code Availability

Sequence data are available from The Global Initiative for Sharing All Influenza Data (GISAID), at https://gisaid.org. The user agreement for GISAID does not permit redistribution of sequences, but lists of the sequences used in our analyses, high-resolution figures, and code will be made available at www.cov.lanl.gov.

### EXPERIMENTAL MODEL AND SUBJECT DETAILS

#### Human Subjects

SARS-CoV-2 sequences were generated using samples taken for routine clinical diagnostic use from 454 individuals presenting with active COVID-19 disease: 243 female, 209 male, 2 no gender specified; ages 18-103 (median 59.6) years.

### METHOD DETAILS

#### Detection and Sequencing of SARS-CoV-2 isolates from clinical samples

Nucleic acid was extracted from 200μl of sample on MagnaPure96 extraction platform (Roche Diagnostics Ltd, Burgess Hill, UK). SARS-CoV-2 RNA was detected using primers and probes targeting the E gene and the RdRp genes for routine clinical diagnostic purposes, with thermocycling and fluorescence detection on ABI Thermal Cycler (Applied Biosystems, Foster City, United States) using previously described primer and probe sets (Corman et al., 2020). Nucleic acid from positive cases underwent long-read whole genome sequencing (Oxford Nanopore Technologies (ONT), Oxford, UK) using the ARTIC network protocol (accessed the 19^th^ of April, https://artic.network/ncov-2019.) Following basecalling, data were demultiplexed using ONT Guppy using a high accuracy model. Reads were filtered based on quality and length (400 to 700bp), then mapped to the Wuhan reference genome and primer sites trimmed. Reads were then downsampled to 200x coverage in each direction. Variants were called using nanopolish (https://github.com/jts/nanopolish) and used to determine changes from the reference. Consensus sequences were constructed using reference and variants called.

#### Data Pipeline

The Global Initiative for Sharing All Influenza Data (GISAID) (Elbe and Buckland-Merrett, 2017; Shu and McCauley, 2017) has been coordinating SARS-CoV-2 genome sequence submissions and making data available for download since early in the pandemic. At time of writing, dozens to hundreds of sequences were being added every day. These sequences result from extraordinary efforts by a wide variety of institutions and individuals: they are an invaluable resource, but are somewhat mixed in quality. The complete sequence download includes a large number of partial sequences, with variable coverage, and extensive ‘N’ runs in many sequences. To assemble a high-quality dataset for mutational analysis, we constructed a data pipeline using off-the-shelf bioinformatic tools and a small amount of custom code.

#### General approach

From theSARS-CoV-2 sequences available from GISAID, we derived a “clean” codon-aligned dataset comprising near-complete viral genomes, without large insertions or deletions (“indels”) or runs of undetermined or ambiguous bases. For convenience in mutation assessment, we generated a codon-based nucleotide multiple sequence alignment, and extracted translations of each reading frame, from which we generated lists of mutations. The cleaning process was in general a process of deletion, with alignment of retained sequences; the following criteria were used to exclude sequences:

1. Fragmented matching (> 20 nt gap in match to reference)
2. Gaps at 5’ or 3’ end (> 3 nt)
3. High numbers of mismatched nucleotides (> 20), ‘N’ or other ambiguous IUPAC codes.
4. Regions with concentrated ambiguity calls: >10 in any 50 nt window)

Any sequence matching any of the above criteria was excluded in its entirety.

#### Sequence mapping and alignment

Sequences were mapped to a reference (bases 266:29674 of GenBank entry NC_045512; i.e., the first base of the orf1ab start codon to the last base of the ORF10 stop codon) using “nucmer” from the MUMmer package (version 3.23; (Kurtz et al., 2004)). The nucmer output “delta” file was parsed directly using custom Perl code to partition sequences into the various exclusion categories (Sequence Mapping Table) and to construct a multiple sequence alignment (MSA). The MSA was refined using code derived from the Los Alamos HIV database “Gene Cutter” tool code base. At this stage, alignment columns comprising an insertion of a single “N” in a single sequence (generating a frame-shift) were deleted, and gaps were shifted to conform with codon boundaries.

Using the initial “good-sequence” alignment, a low-effort parsimony tree was constructed using PAUP: a single replicate heuristic search using stepwise random sequence addition. Sequences in the alignment were sorted vertically to correspond to the (ladderized) tree, and reference-sequence reading frames were added.

#### Data partitioning and phylogenetic trees

Estimated phylogenies were inferred for three distinct data partitions: the full sequence set (near-complete genomes), the spike open reading frame, and the full set with the spike open reading frame (orf) excluded. The full genome tree was used for Fig. 1. The tree with the spike orf excluded was intended to allow independent assessment of the phylogenetic distribution of changes within the spike protein, by preventing convergent or homoplastic mutations driven by phenotypic selection upon the spike protein from overwhelming phylogenetic signal from the rest of the genome. We confirmed that the phylogenetic observation discussed in Fig. 1 were supported in the Spike-excluded tree, but do not include it here. It is available for cross-checking phylogenetic based inferences at www.cov.lanl.gov.

Trees were inferred by either of two methods: 1. neighbor-joining using a p-distance criterion, (Swofford, 2003) or 2. parsimony heuristic search using a version of the parsimony ratchet (Goloboff, 2014).

#### Global Maps

The Covid-19 pie chart map is generated by overlaying Leaflet (a JavaScript library for interactive maps) pie charts on maps provided by OpenStreetMap. The interface is presented using rocker/shiny, a Docker for Shiny Server.

#### Sequence quality control identification of a sequencing error

We discovered a sequencing processing error that gave rise to what appeared to be a mutation at position 943 (24389 A>C and 24390 C>G) in Spike that was evident in sequences from Belgium (Fig. S7). We contacted the group in Belgium, the source of the data, who were already aware of the issue, concurred with our interpretation, and they had been in touch with GISAID with a request to remove the problematic sequences. The error was not found among more recent sequences from Belgium (Fig. S6).

We identified the issue with this site as part of another study using a method to detect systematic sequencing errors (Freeman et al., 2020); we are interrogating the quality of available sequencing data and these positions were highlighted as suspect. We interrogated these positions in the raw sequencing data from Sheffield, and although these two variants are not present in the final consensus sequence from any of the Sheffield isolates, the raw, untrimmed bam files show their presence in only one of the amplicons covering the site (Fig. S8 A&B). We noticed that in fact this position is to the left of the 5’ primer of amplicon 81 in what we believe to be an adapter sequence. Comparison of the WuHan reference and the adapter sequence reveals similarity around this position:

Nanopore adapter sequence: CAGCACCTT
The WuHan reference sequence: CAGCAAGTT

In our validation set, we see a C present at around 50% of called bases at both these positions in raw data but this region is trimmed by the ARTIC pipeline and is therefore not used to call variants and contribute to the final consensus sequence. Although it is evident in amplicon 81, in this region, there is no evidence for these variants in the data from amplicon 80, which also covers these positions. We include a figure (Fig. S8) that hopefully will help to explain our finding.

In summary this is an error that has arisen due to a combination of improper trimming of adapter and primer regions from raw sequencing reads before downstream analysis, and the coincidental homology between the nanopore adapter sequence and the WuHan reference genome in this region.

### QUANTIFICATION AND STATISTICAL ANALYSIS

#### Clinical data and disease severity

To assess possible associations of clinical and sequence variables with disease severity, we used a Generalized Linear Model (GLM) using outpatient vs. hospitalized status as outcome and age, gender, D614 and PCR cycle threshold for E gene amplification (E_gene_CT) as potential predictors. Outpatient vs. hospitalized status, gender, and D614 were all categorized as binomial factors, while age and E_gene_CT were considered as continuous variables. We started with the largest model that included all variables and then used ANOVA to down-select the best predicting model. All coding was done using R and the lme4 package (Bates 2015).

#### Mutation rate analysis of Belgian sequences

The phylogenetic tree was broken up into subtrees using the R ape package (Paradis and Schliep 2018), and those subtrees containing only Belgian or only non-Belgian tips were selected, and their total branch lengths calculated. For each subtree the R package phangorn (Schliep 2011) was used to calculate the minimum number of changes required at each site. The maximum rate, in mutations per unit branch length, compatible with the non-Belgian data was calculated as its 95% upper confidence level assuming a Poisson distribution of mutations with the Poisson parameter proportional to the branch length. The p-value of the Belgian data was estimated from a Poisson distribution with parameter given by the rate for non-Belgian data multiplied by the total branch length of the Belgian trees. Only two sites were found to be significant after Bonferroni corrected by the number of sites in the alignment.

#### Identification of recombination candidates using RAPR

To identify the candidate recombination parent and child sequences shown in Fig. S8, we first did all triplet comparisons of all sequences from local region using the RAPR, and the raw p-values based on a run-length statistic all the comparisons were rank-ordered to identify the triplet candidates with strongest evidence for recombination to be used as a basis for further exploration. Thus these p-values are uncorrected for multiple testing and not formally compared against the alternative hypotheses of stepwise convergent mutation, as we traditionally do in RAPR analysis (Song et al., 2018). However, given the very low overall mutation rate among SARS-CoV-2 pandemic, it is extremely unlikely that the mutational patterns seen in the recombinant sequences were a result of stepwise convergence.

### ADDITIONAL RESOURCES

Archived data for the current manuscript, and current data updates, analytical results, and webtools: https://COV.lanl.gov.

### KEY RESOURCES TABLE

The R Foundation for Statistical Computing, http://www.R-project.org

**R packages (https://cran.r-project.org/ except as noted)**

- The R Foundation for Statistical Computing, http://www.R-project.org

R packages (https://cran.r-project.org/ except as noted)

- phangorn (version 2.5.5)
- ggplot2 (version 3.3.0)
- beeswarm (version 0.2.0)
- tidyverse (version 1.3.0)
- ape (version 5.3)
- lme4 (version 1.1.21)
- data.table (version 1.12.8): https://github.com/Rdatatable/data.table

**Web tools, software, and protocols**

ADD COVID tools

- GISAID: https://www.gisaid.org
- Los Alamos SARS-CoV-2 mutation analysis pipeline: https://COV.lanl.gov
- RAPR: Recombination Analysis PRogram. https://www.hiv.lanl.gov/content/sequence/RAP2017/rap.html
- VMD: Visual Molecular Dynamics: https://www.ks.uiuc.edu/Research/vmd/
- Highlighter: A tool to highlight matches, mismatches, and specific mutations in aligned protein or nucleotide sequences. https://www.hiv.lanl.gov/content/sequence/HIGHLIGHT/highlighter_top.html
- Rainbow Tree: https://www.hiv.lanl.gov/content/sequence/RAINBOWTREE/rainbowtree.html
- Aliview, a sequence alignment viewer and editor: https://ormbunkar.se/aliview/
- Add COVID tools
- Sequence resconstruction: ARTIC network protocol (accessed the 19^th^ of April, https://artic.network/ncov-2019.)
- Base call resolution: Nanopolish (https://github.com/jts/nanopolish)
- Matplotlib: A 2D Graphics Environment (Hunter, 2007)
- PAUP https://paup.phylosolutions.com

