## Supplemental Information for "Spike mutation pipeline reveals the emergence of a more transmissible form of SARS-CoV-2"

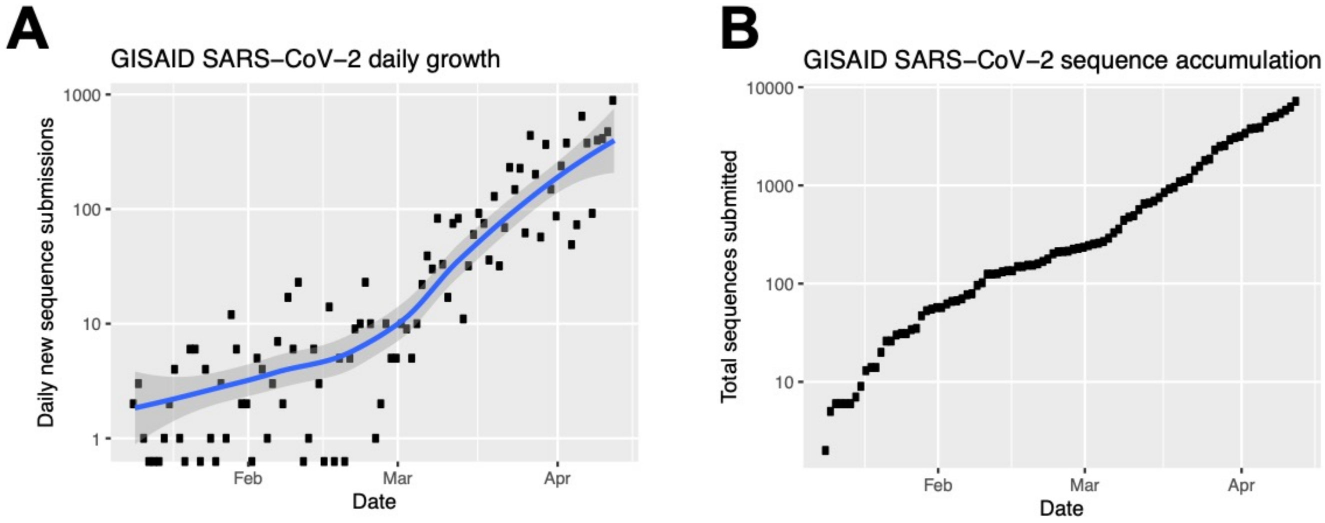


**Fig. S1. The submission of new SARS CoV-2 sequences to the global initiative on sharing avian flu data (GISAID) each day.** The number of sequences submitted each day to GISAID is increasing at a rapid pace, and the cumulative number of sequences has reached nearly 10,000 sequences in the 4 months since the epidemic began. We end here at April 13, the point at which we did the bulk of the analysis in this paper, but our goal is a daily update of the figures such as the trees that track accruing Spike mutations shown in Fig. 1.

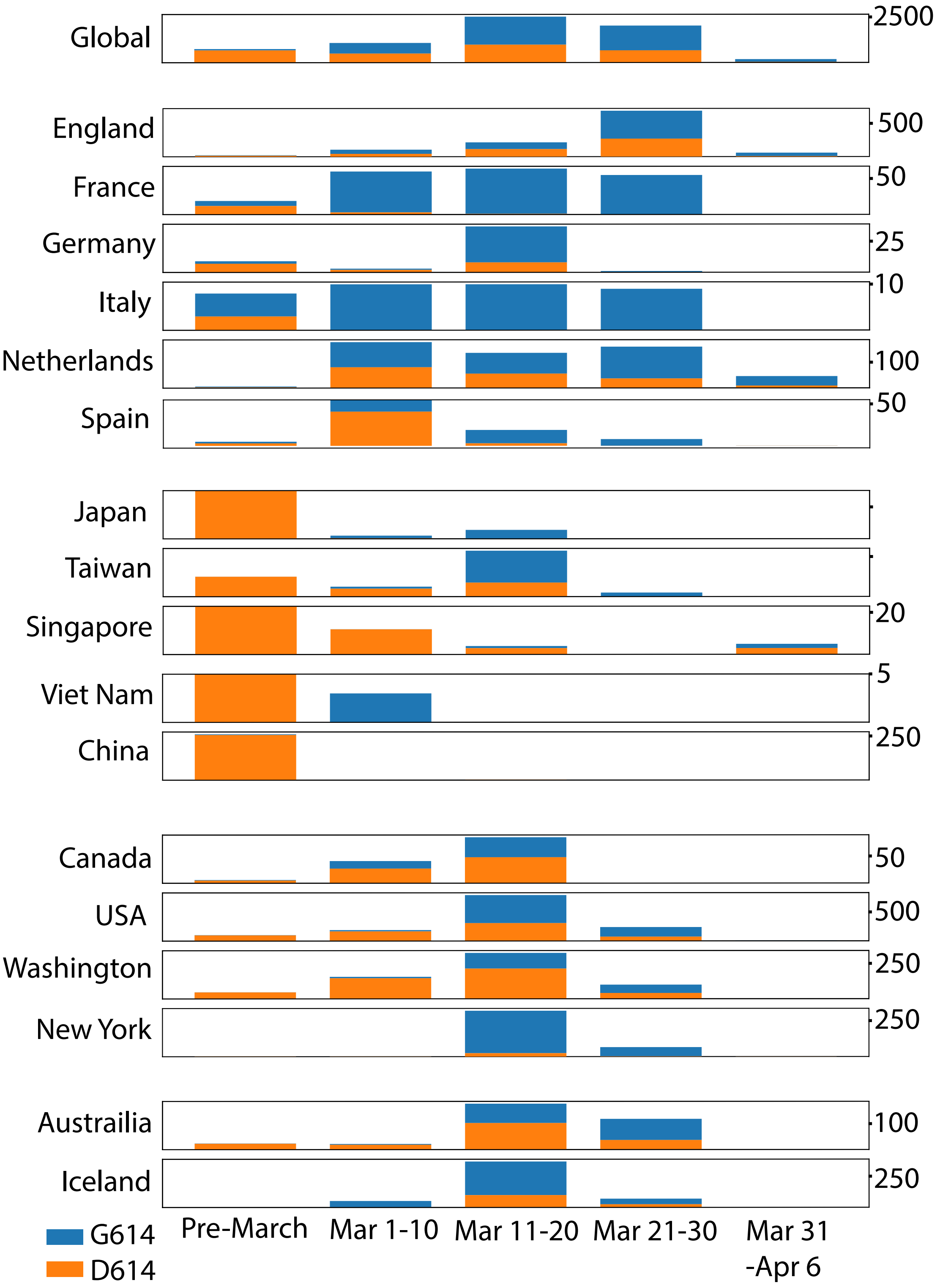


**Fig. S2. The increasing frequency of the D614G mutation over time, in different regions of the world.** The figure complements Fig. 2B in the main text, but here the height of the bars indicates the tally of the number of sequences of each form, rather than the frequency as in Fig. 2B.

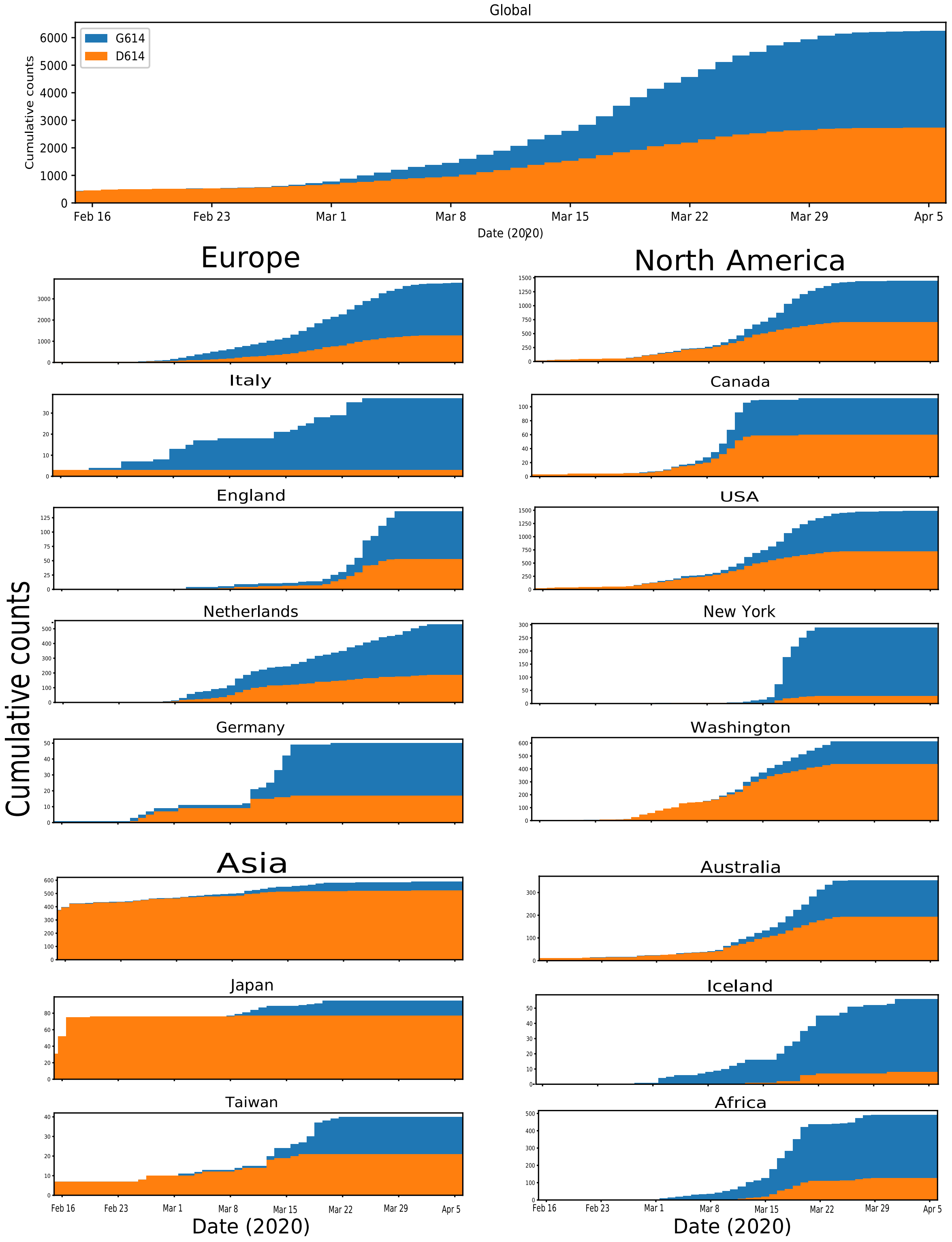


**Fig. S3. Daily cumulative plots showing the relative amount of D614 (orange) and G614, (blue) in different regions of the world**. These plots present the same data as in Fig. 3 in a different view, shown as cumulative daily tallies rather than the running weekly averages shown in Fig. 3

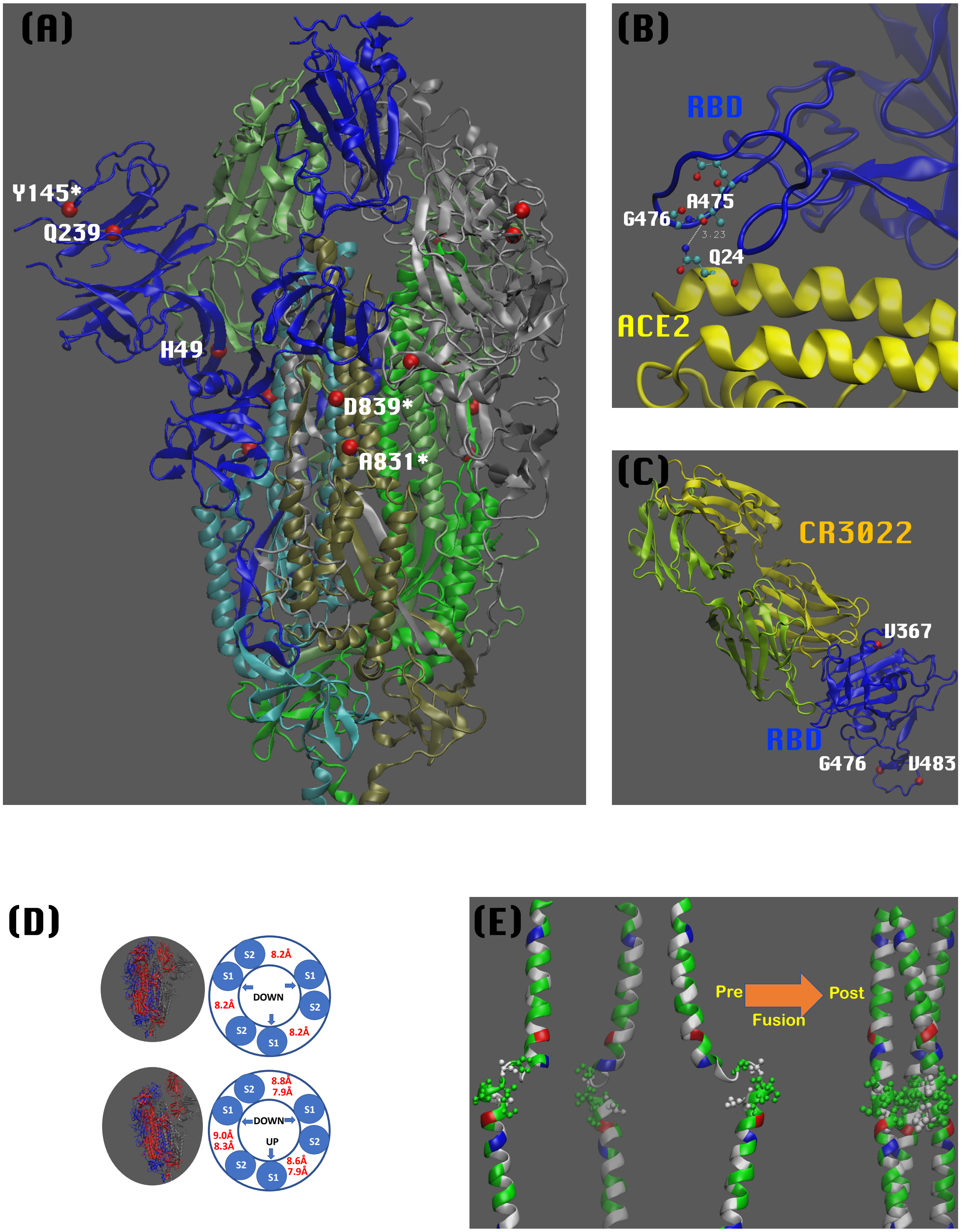


**Fig. S4. Additional structural mapping relating to sites of interest in the Spike protein.**
**A)** Structural mapping of additional mutational sites in the Spike protein. (PDB:6VSB). Different colors are used to distinguish the protomers and S1 and S2 units (protomer #1: S1-blue, S2-cyan; protomer #2: S1-grey, S2-tan; protomer #3: S1-light green, S2-dark green). The RBD of

protomer #1 is in “UP” position for engagement with ACE2 receptor. Red balls are used to indicate individual mutational sites. When a mutational site is missing in the structure, the closest residue to the mutational site is mapped and is indicated by a * next to the amino acid label. **B)** The RBD-ACE2 binding interface (PDB: 6M17) near the mutational site G476S. Q24 in ACE2 is the closest residue to this mutational site. However, the neighbor residue, A475, is even closer to the ACE2 with a potential to form hydrogen bonding. **C)** The three mutational sites in RBD in the context of recognition by the antibody CR3022 (PDB: 6W41). The heavy and light chains are colored in yellow and green, respectively. **D)** Structure based studies imply that the RBD needs to be in the “UP” position to engage with ACE2 receptor. We evaluate whether the protomer-protomer contacts between D416 and T859 could be impacted by the “UP” or “DOWN” conformations. Here we show the protomer-protomer distance (red) defined by the CA atoms of these residues in a cartoon diagram representing all-DOWN and one-UP configurations of RBD in a Spike trimer. In the all-DOWN conformation (PDB: 6VXX), the inter-protomer distance between *D614* and T859 does not change within the trimer. In the one-UP conformation, inter-protomer distances slightly vary within a trimer. In the case of one-UP conformation, two distances are provided. The top (PDB: 6VSB) and bottom (PDB: 6VYB) numbers are extracted from different cryo-EM structures. **E)** The left and right images show the amphipathicity of pre-fusion (PDB:6VSB) and post-fusion (PDB: 6LXT) helices of HR1 region (Charged/Polar: red/blue/green and hydrophobic: white). Ser/Thr rich region spanning *S937-S943* (ball and sticks) may play a role in elongation and association of these amphipathic helices. This figure accompanies Fig. 4 in the main text.

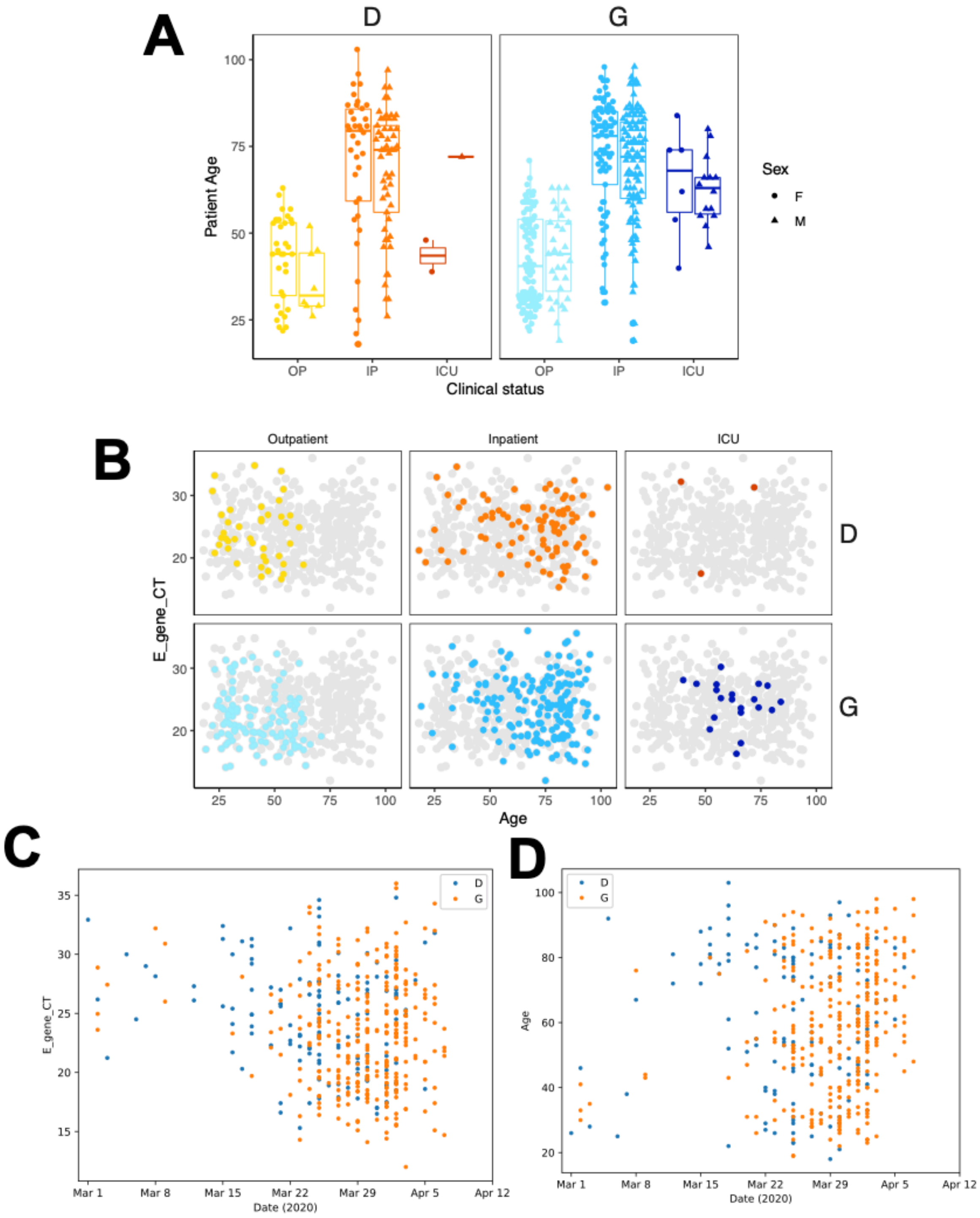


**Fig. S5. Distribution of clinical status with respect to patient age, sex, and a proxy for viral load when sampled. A)** Patient age and sex by clinical status and D614G status. OP = outpatient, IP = inpatient, ICU = intensive care unit. **B)** Cycle threshold for product detection in PCR of the E gene: lower values correspond to greater quantities of viral RNA. In each panel, colored points denote data from that panel; grey points, representing the total data set, are identical for all 6 panels. Colors correspond to those in the main manuscript (greater intensity reflecting more severe disease as indicated by OP, IP or ICU care decisions. The relative distribution over sampling time of D614 and G614 among values for **C)** PCR CT, and **D**) Age. The relative distribution of D614 and G614 appears to be reasonable consistent though time, indicating that there was not an overt bias introduced due to G614 tending to be sampled later and D614 sampled earlier. Of note, the earliest samples have high E gene CT values, indicative of lower viral loads, and the patient population tends to be younger. This figure accompanies Fig. 5 in the main text.

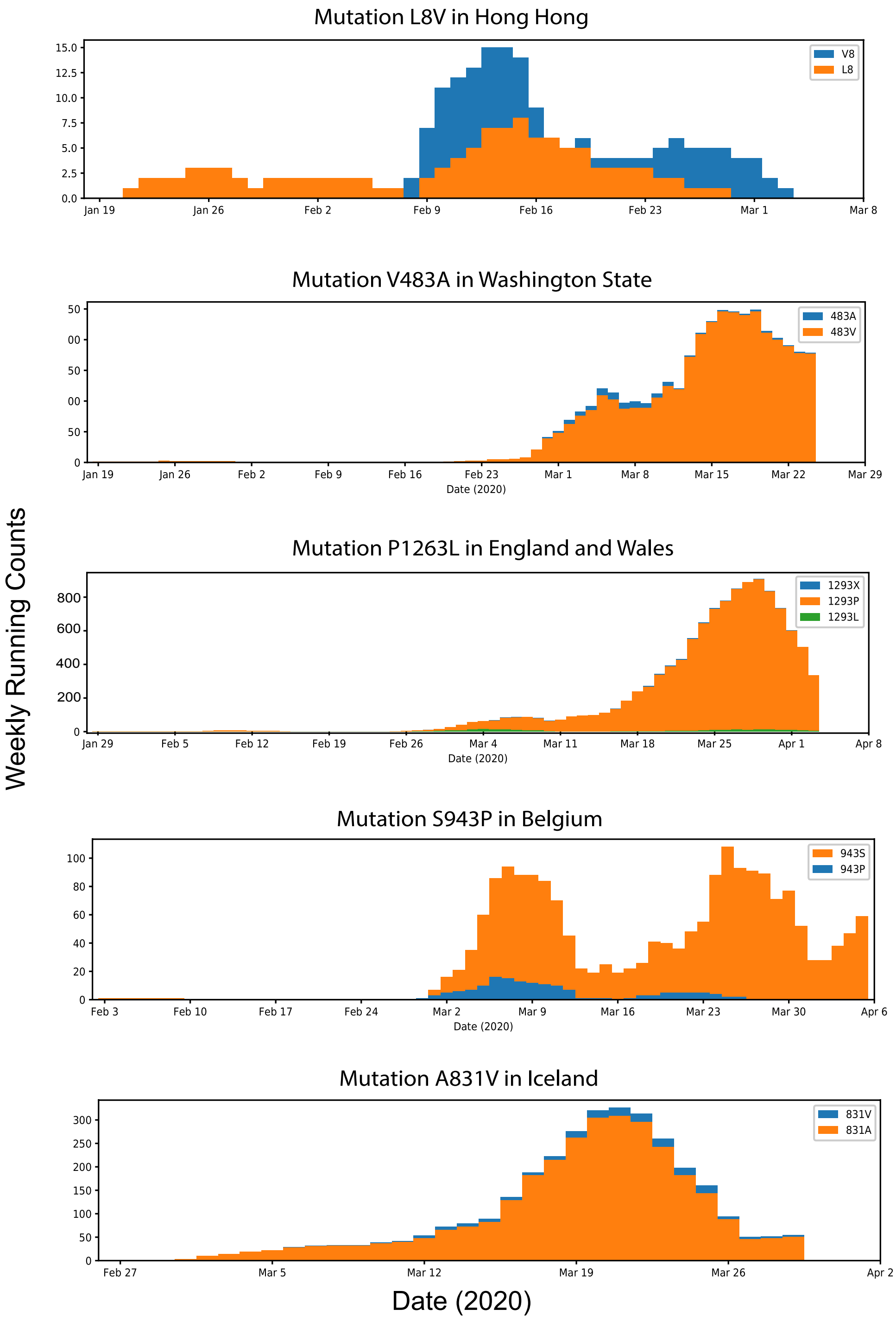


**Fig. S6. Weekly running counts of mutations in sites of interest in local epidemics where a particular mutation of interest was enriched.** This figure accompanies Table 1 in the main text.

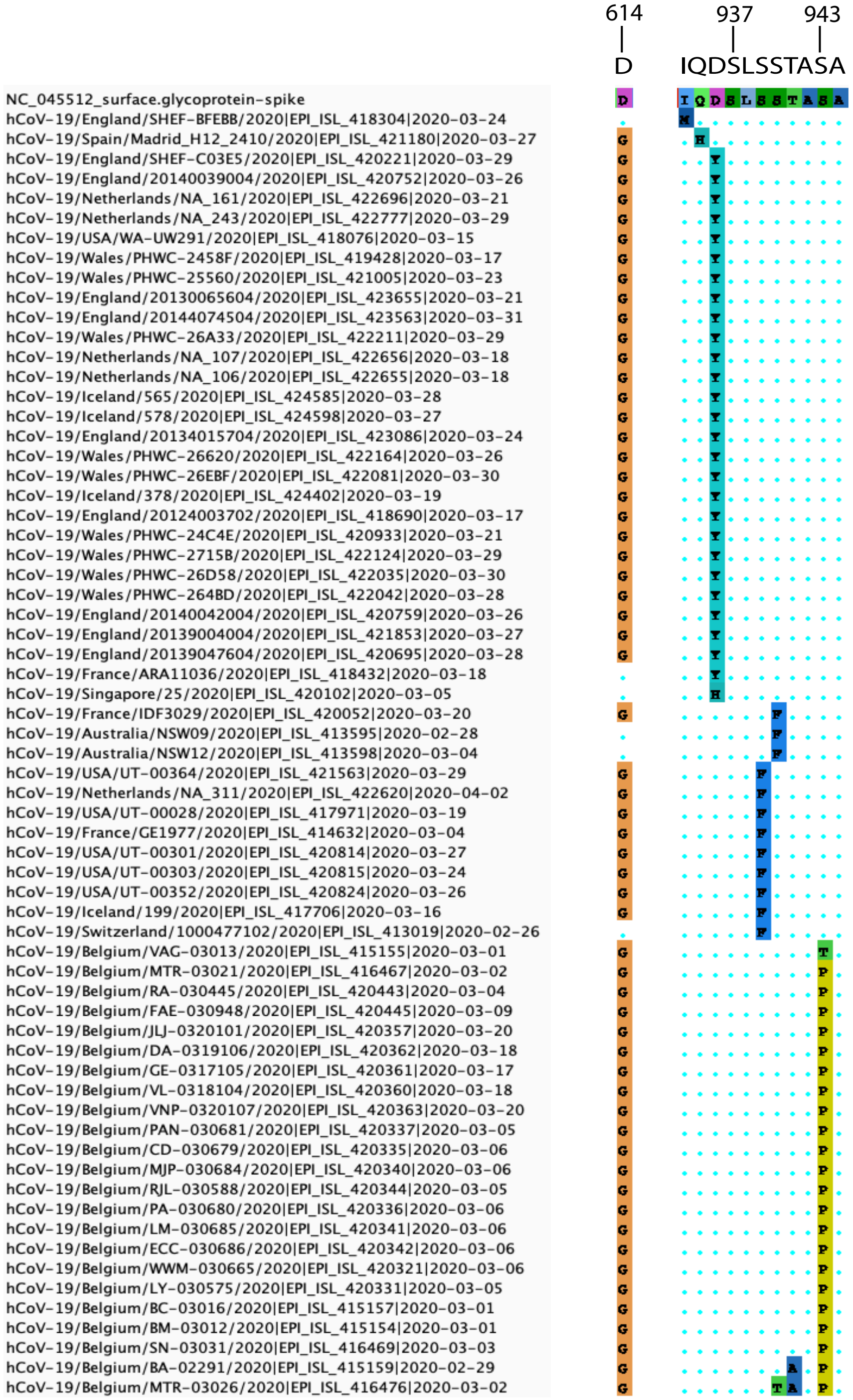


**Fig. S7.** This figure shows a cluster of mutations that give rise a entropy region when scanning the Spike protein. It contains a Serine rich motif may be important for helical formation and fusion. The D936Y mutation are beginning to be observed both within and outside of the major clade defined by the D614G mutation (shown to the left of the region of interest) and in many diverse regions geographic regions, suggesting de novo mutations or local recombination events, as are the S939F and S940F changes. The S943P mutation was shown to be a sequencing artifact. This figure accompanies Fig. 4 in the main text.

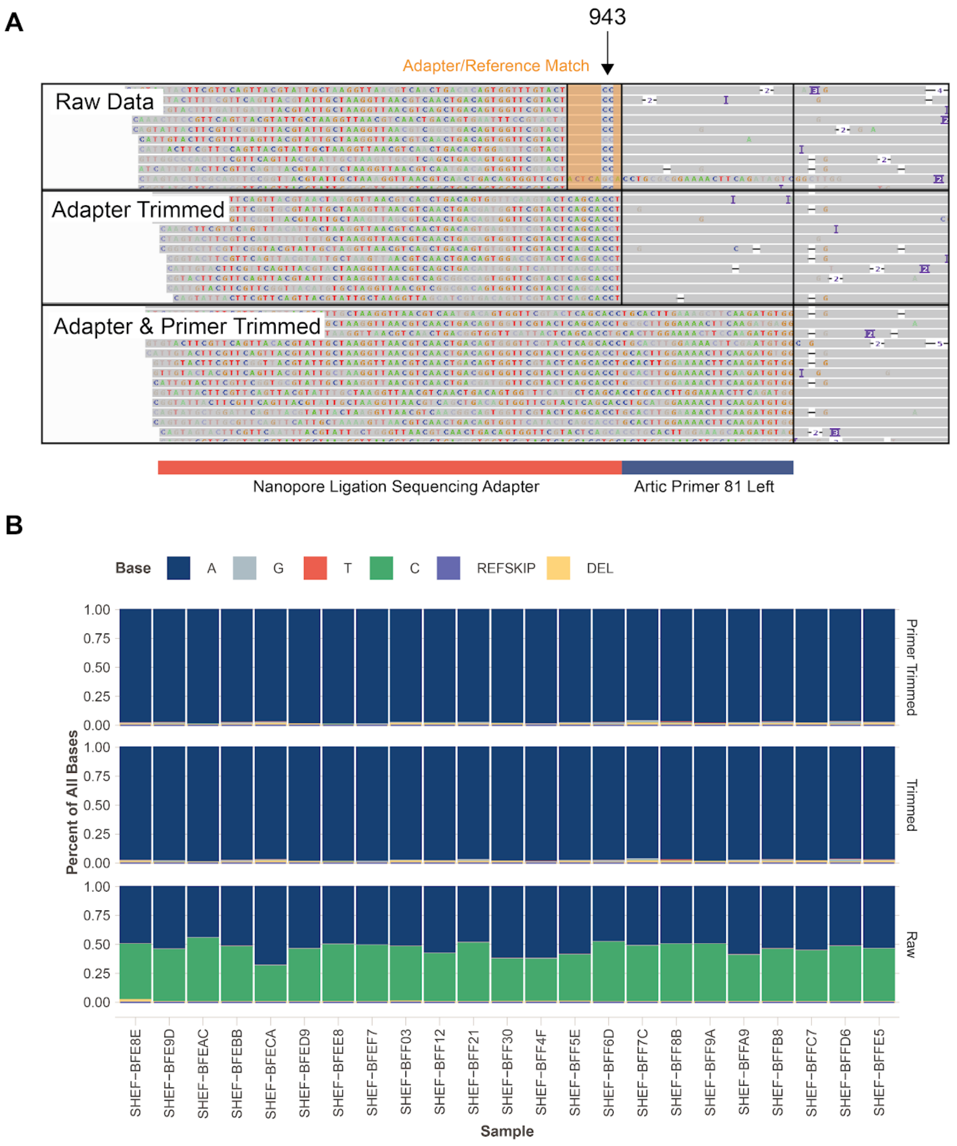


**Fig. S8. Investigation of S943P. A.** IGV plots showing bam files from nanopore sequencing data of amplicons produced by the Artic network protocol. Raw data from amplicon 81 contains a portion of adapter sequence which is homologous to the reference genome, apart from the C variants which lead to a S943P mutation call. This region is therefore included in variant calling if location-based trimming is not carried out. Subsequent panels show that this region is soft clipped when trimming adapters and primers and is therefore not available for variant calling. **B.** Base frequencies at position 24389 in 23 samples from the Sheffield data show that C is present in half of the reads in the raw data, but is absent from trimmed and primer trimmed data. This figure is associated with the Star Methods.
